# Quantitative Analysis of Coupled Carbon and Energy Metabolism for Lignin Carbon Utilization in *Pseudomonas putida*

**DOI:** 10.1101/2025.03.24.645021

**Authors:** Nanqing Zhou, Rebecca A. Wilkes, Xinyu Chen, Kelly P. Teitel, James A. Belgrave, Gregg T. Beckham, Allison Z. Werner, Yanbao Yu, Ludmilla Aristilde

## Abstract

Soil *Pseudomonas* species, which can thrive on lignin-derived phenolic compounds, are widely explored for biotechnology applications. Yet, there is limited understanding of how the native metabolism coordinates phenolic carbon processing with cofactor generation. Here, we achieve quantitative understanding of this metabolic balance through a multi-omics investigation of *Pseudomonas putida* KT2440 grown on four common phenolic substrates: ferulate, *p-*coumarate, vanillate, and 4-hydroxybenzoate. Relative to succinate as a non-aromatic reference, proteomics data reveal >140-fold increase in proteins for transport and initial catabolism of each phenolic substrate, but metabolomics profiling reveals that bottleneck nodes in initial phenolic compound catabolism maintain more favorable cellular energy state. Up to 30-fold increase in pyruvate carboxylase and glyoxylate shunt proteins implies a metabolic remodeling confirmed by kinetic ^13^C-metabolomics. Quantitative analysis by ^13^C-fluxomics demonstrates coupling of this remodeling with cofactor production. Specifically, anaplerotic carbon recycling via pyruvate carboxylase promotes fluxes in the tricarboxylic acid cycle to provide 50-60% NADPH yield and 60-80% NADH yield, resulting in 2-fold higher ATP yield than for succinate metabolism; the glyoxylate shunt sustains cataplerotic flux through malic enzyme for the remaining NADPH yield. The quantitative blueprint elucidated here explains deficient versus sufficient cofactor rebalancing during manipulations of key metabolic nodes in lignin valorization.

---

Lignin, a structural polymer in plant biomass that is considered the second most abundant biopolymer after cellulose, represents an important renewable carbon source^1^. Chemical depolymerization of lignin yields various phenolic compounds, which can be used directly as platform chemicals or as feedstocks for bioproduction^2–8^. Lignin-related phenolic structures include hydroxybenzoates (4-hydroxybenzoate, vanillate, gallate, and syringate) and hydroxycinnamates (*p*-coumarate and ferulate)^9^. Soil *Pseudomonas* species have native metabolic capabilities to process benzoate, 4-hydroxybenzoate (4HB), vanillate (VAN), gallate, *p*-coumarate (COU), and ferulate (FER)^10–18^. Of particular interest is *Pseudomonas putida* KT2440, which is extensively explored as a chassis for biotechnology^19–22^. Encoded in *P. putida* KT2440 are pathways for the catabolism of hydroxybenzoates and hydroxycinnamates involving *ortho*-cleavage of the intermediate protocatechuate (PCA) through β-ketoadipate to generate carbon influx into the tricarboxylic acid (TCA) cycle in the central carbon metabolism^13^ (Fig. 1A). In the peripheral pathways upstream of β-ketoadipate, the uptake and catabolism of hydroxycinnamates (COU and FER) and hydroxybenzoates (4HB and VAN) to PCA are catalyzed by a series of enzymes with distinct cofactor specificities^13,23–28^ (Fig. 1A). Previous metabolic engineering of *P. putida* targeted directing carbon influx from phenolic carbons from initial catabolism towards desired products such as PCA, vanillin, β-ketoadipate, muconic acid, pyridine 2,4-dicarboxylic acid, adipic acid, indigoidine, and polyhydroxyalkanoates^29–42^. Metabolic bottlenecks, which were inferred from extracellular accumulation of VAN in FER-fed cells or of 4HB in COU-fed cells^33,40,41,43–45^, could not be overcome completely via genetic engineering due to cofactor deficiency^33,43,41^. Therefore, metabolic engineering to enhance phenolic carbon conversion must address appropriate cofactor supply. However, largely lacking is a quantitative understanding of how the native metabolic network in *P. putida* KT2440 couples carbon fluxes from phenolic acid structures with cofactor production.

**Fig. 1.**
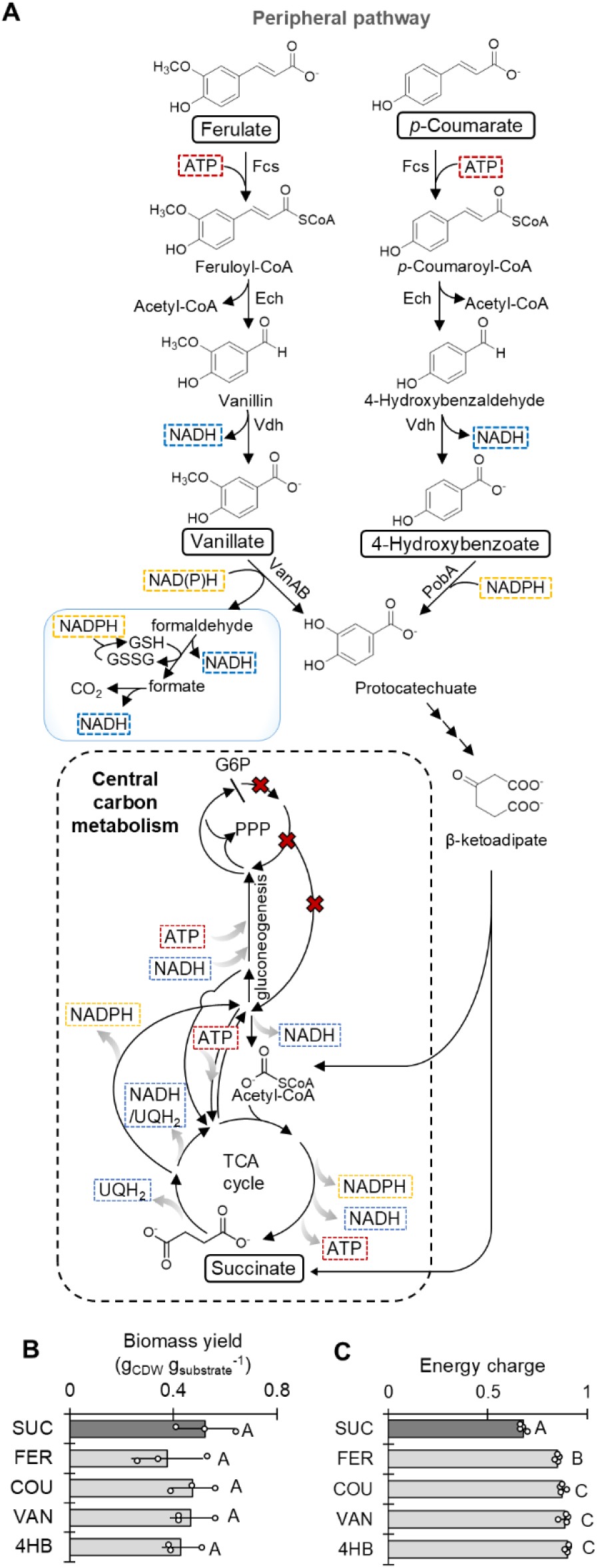
Favorable energy charge from the metabolism of phenolic acid structures. (A) Schematic illustration of cofactor investment and production in the peripheral pathway and central carbon metabolism. (B) Biomass yield of *P. putida* cells utilizing SUC, FER, COU, VAN, and 4HB. Data are expressed as mean ± standard deviation from three biological replicates (*n* = 3). (C) Energy charge calculated from quantified ATP, ADP, and AMP pools when *P. putida* cells were fed with SUC, FER, COU, VAN, and 4HB. Energy charge = ([ATP] + 0.5 × [ADP]) / ([ATP] + [ADP] + [AMP]). Data are expressed as mean ± standard deviation from four independent biological replicates (*n* = 4). In B and C, data for SUC-fed condition were adapted from Wilkes et al.^60^ Abbreviations: GSH, glutathione; GSSG, glutathione disulfide. The red crosses represent no carbon flux through the oxPPP and Entner–Doudoroff pathway during gluconeogenesis. In B and C, one-way analysis of variance (ANOVA) was performed followed by Tukey’s HSD post hoc test. Statistically significant differences (*P* < 0.05) are denoted by a change in letter.

Metabolic flux modeling or fluxomics constrained by ^13^C-metabolomics data provides quantitative analysis of carbon fluxes^46–53^. Previous ^13^C-fluxomics studies of *P. putida* focused on glycolytic metabolism for sugars (i.e., glucose, xylose) or gluconeogenic metabolism for succinate (SUC)^18,54–63^. The only two fluxomics studies of aromatic carbon metabolism in *P. putida*, for benzoate alone or with glucose as a co-substrate^17,18^, both illustrated the activation of the glyoxylate shunt, which conserved carbon by channeling isocitrate away from the canonical decarboxylation steps in the TCA cycle. Flux through the glyoxylate shunt can feed into malic enzyme for NADPH generation, or malate dehydrogenase, for NADH or ubiquinol generation. Interestingly, the flux through malate dehydrogenase was either 3-fold higher than the flux through malic enzyme or there was only flux through malate dehydrogenase, both of which would be advantageous to satisfy the NADH demand for benzoate catabolism through the catechol pathway in *P. putida*^17,18^. For the catabolism of hydroxybenzoate and hydroxycinnamate structures, which is funneled through PCA instead of catechol, it remains unknown how metabolic fluxes would be partitioned in *P. putida* to optimize the catabolic pathways with cofactor supply (Fig. 1A).

While ATP in aerobic bacteria is generated primarily through oxidative phosphorylation of NADH produced in glycolysis and the TCA cycle^64,65^, the supply of NADPH required for biosynthesis and stress tolerance is produced from different metabolic routes. During growth on glucose, NADPH production in *P. putida* is mainly through glucose-6-phosphate dehydrogenase and 6-phosphogluconate dehydrogenase in the oxidative pentose phosphate pathway (oxPPP)^56,59^. However, minimal to no flux in oxPPP was reported during growth on two gluconeogenic substrates (SUC or benzoate)^17,60^. Instead, SUC metabolism in *P. putida* relied on high flux through isocitrate dehydrogenase and malic enzyme in the TCA cycle to produce NADPH, and on transhydrogenase reactions to replenish NADPH from excess NADH pool^60^. Whether *P. putida* would rely on similar metabolic routing or energetic bypass for NADPH production during phenolic substrate utilization remains to be elucidated.

The cataplerotic and anaplerotic nodes, which distribute respectively carbon fluxes from the TCA cycle to gluconeogenesis and from lower glycolysis to the TCA cycle^66^, have implications for cofactor balance (Fig. 1A). In the cataplerotic direction, due to lack of phosphoenolpyruvate (PEP) carboxykinase for the decarboxylation of oxaloacetate (OAA) to PEP^67^, *P. putida* KT2440 must rely on either malic enzyme to convert malate to pyruvate with NADPH production or OAA decarboxylase to convert OAA to pyruvate with no cofactor involved^68^. In the anaplerotic direction, pyruvate carboxylation to OAA consumes ATP, while PEP carboxylation to OAA has no cofactor input^68^. It remains unclear how *P. putida* KT2440 would modulate carbon flux distribution between cataplerosis and anaplerosis to meet the demand for reducing equivalents and energy required for phenolic carbon catabolism.

Here, our central hypothesis was that remodeling of carbon flux into and within the TCA cycle by *P. putida* KT2440 would promote the net production of NAD(P)H and ATP to satisfy the cofactor demands in the *p*-coumaroyl and coniferyl pathways (Fig. 1A). We tested our hypothesis by employing genetic engineering, targeted metabolomics, whole-cell proteomics, kinetic ^13^C-metabolomics, and ^13^C-fluxomics during feeding of *P. putida* KT2440 on FER, COU, VAN, or 4HB. First, using quantitative metabolomics for cofactor profiling in wild-type and engineered strains, we investigated how the cellular energy state would be influenced by alterations in the aromatic carbon influx in several strains that overexpressed key bottleneck-relevant genes. Second, to construct the metabolic network that establishes the cellular energy state, we profiled protein levels to identify metabolic remodeling nodes and verify these nodes using kinetic ^13^C-profiling in metabolites coupled with ^13^C-carbon mapping. Third, we integrated the proteomics and ^13^C-metabolomics data to perform ^13^C-fluxomics to map quantitatively carbon fluxes through the metabolic network. Fourth, we leveraged the quantitative flux analysis to determine production and consumption rates of cofactors, thereby evaluating metabolic flux controls on cofactor maintenance during conversion of the different lignin-derived phenolic compounds. Our findings provide quantitative insights into the relationship between carbon metabolism and energy metabolism in *P. putida* KT2440 utilizing different hydroxycinnamate and hydroxybenzoate substrates related to lignin.

## Results

### Bottleneck Nodes in Initial Phenolic Substrate Catabolism Sustain Favorable Cellular Energy Balance

During growth on 100 mM C of FER, COU, VAN or 4HB as the sole carbon source, *P. putida* KT2440 showed growth rates ranging from 0.53-0.88 h^-1^ [Supplementary Information (SI), Fig. S1A], which were 10-41% slower compared to growth rate reported for SUC^60^, a non-aromatic substrate reference. Substrate depletion rates were comparable for SUC and 4HB (16.0 ± 2.0 and 15.9 ± 3.1 mmol g_CDW_^-1^ h^-1^, respectively) (*P =* 0.96), but the corresponding rates for FER, COU, and VAN were 58%, 38%, and 49% lower than SUC, respectively (*P* < 0.05) (SI, Fig. S1B). Despite the differences in growth and substrate depletion rates, the biomass yield of *P. putida* was similar across all five substrates (*P* = 0.64-0.99) (Fig. 1B). Since biomass synthesis requires energy input, we further evaluated the energy status of *P. putida* during feeding on the different substrates by calculating the energy charge using quantified intracellular levels of ATP, ADP, and AMP^60^ (SI, Table S1). Remarkably, we obtained 25-31% higher energy charge when cells were fed on the phenolic substrates compared to SUC feeding (*P* < 0.001) (Fig. 1C; SI, Table S1). These data implied that carbon influx during phenolic acid assimilation was optimized for favorable cellular energy charge.

With respect to regulation of carbon influx, bottlenecks were proposed at three nodes in initial phenolic acid catabolism, based on extracellular metabolic overflow in exponentially growing cells: the VanAB node in the coniferyl branch^33,40^, the PobA node in the *p*-coumaroyl branch^33,41,43^, and the PcaHG node downstream of both branches^40^ (Fig. 1A). Here, we obtained direct evidence using intracellular metabolomics analysis and ^13^C kinetic isotopic profiling of exponentially growing cells to confirm the three reported bottlenecks, in addition to identifying one at the Vdh node (Fig. 2). We evaluated changes in cellular energy charge due to resolving the bottlenecks with gene overexpression (Fig. 2).

**Fig. 2.**
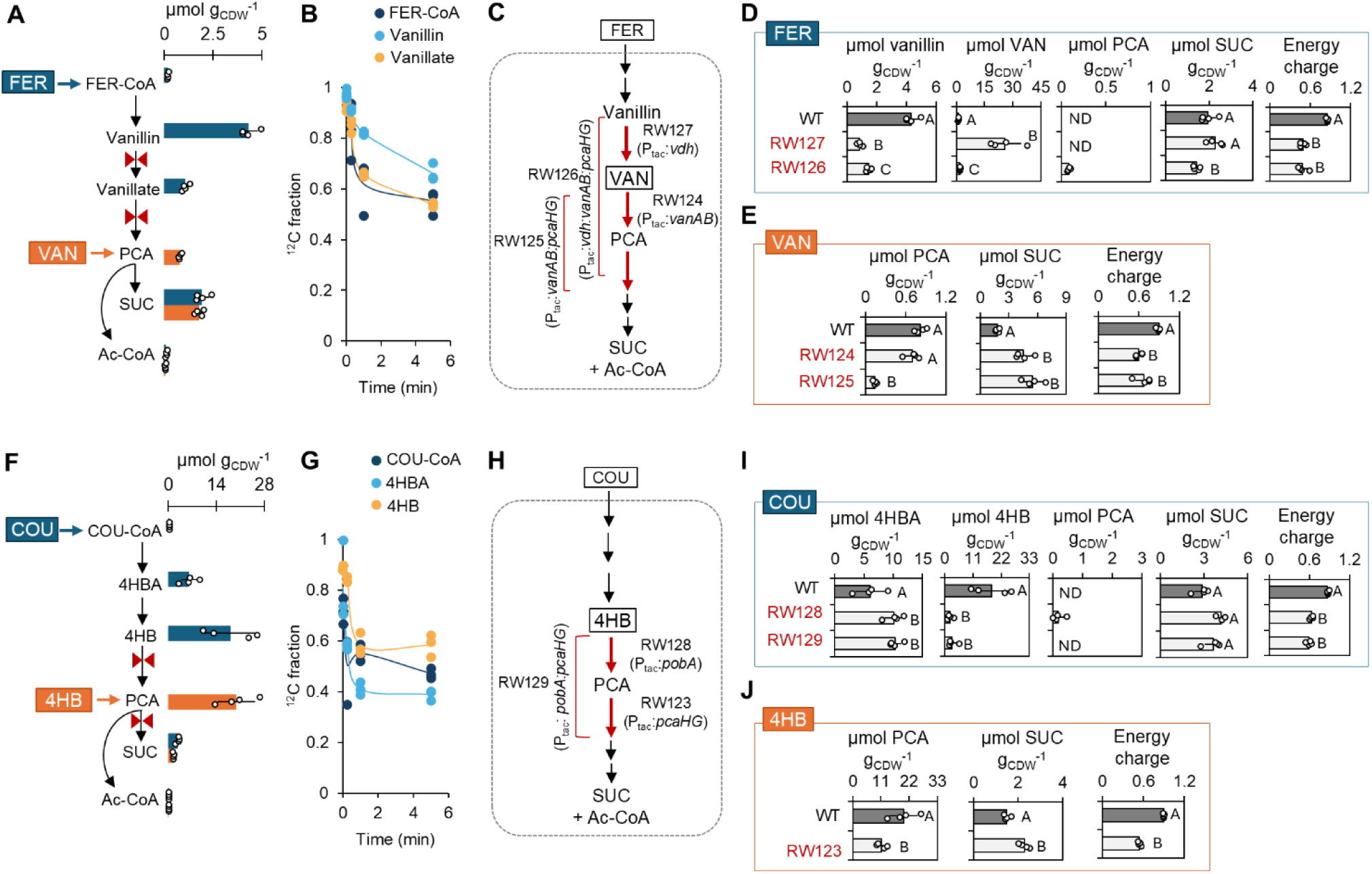
Intracellular evidence of bottleneck nodes in the peripheral pathways of phenolic substrate catabolism. (A) Intracellular levels of intermediates involved in FER and VAN utilization. (B) Kinetic depletion in unlabeled (^12^C) fraction of key metabolites in the coniferyl branch during incorporation of ^13^C-FER. (C) Targeted nodes in the coniferyl branch for gene overexpression. Intracellular metabolite levels and energy charge in the wild-type and mutant strains during feeding on (D) FER and (E) VAN. (F) Intracellular levels of intermediates involved in COU and 4HB utilization by *P. putida.* (G) Kinetic depletion in the unlabeled (^12^C) fraction of key metabolites in the *p*-coumaroyl branch during incorporation of ^13^C-COU. (H) Targeted nodes in the *p*-coumaroyl branch for gene overexpression. Intracellular metabolite levels and energy charge in the wild-type and mutant strains during feeding on (I) COU and (J) 4HB. In B and F, red bowties indicate the identified bottlenecks. ND: not detected. Abbreviations: FER-CoA, feruloyl-CoA; COU-CoA, *p*-coumaroyl-CoA. In D, E, and I, one-way ANOVA was performed followed by Tukey’s HSD post hoc test to determine the significance. In J, unpaired *t*-test was performed to evaluate the significance. Statistically significant differences (*P* < 0.05) are denoted by a change in letter. All data in A, D, E, F, I, and J are obtained from four independent biological replicates (*n* = 4). All data in B and G were obtained from three independent biological replicates (*n* = 3).

In the coniferyl branch, bottleneck at the Vdh node was characterized by vanillin accumulation (4.3 ± 0.5 µmol g_CDW_^-1^) relative to both the precursor feruloyl-CoA (0.2 ± 0.03 µmol g_CDW_^-1^) and the downstream metabolites vanillate (1.1 ± 0.2 µmol g_CDW_^-1^) and PCA (below the detection limit) (Fig. 2A). This bottleneck was further illustrated by 50% lower incorporation of ^13^C-FER in vanillin compared to feruloyl-CoA within 1 min of isotope switch (*P* < 0.05) (Fig. 2B). Overexpression of *vdh* (strain RW127) led to an 80% decrease in vanillin and a 24-fold increase in VAN (*P* < 0.001), indicating that addressing the bottleneck at Vdh triggered a downstream bottleneck at VanAB (Fig. 2C). Even with VAN as the direct carbon source, PCA level remained low (0.8 ± 0.1 µmol g_CDW_^-1^), due to inefficient VAN conversion by VanAB (Fig. 2C). In relation to this bottleneck, we constructed three strains: RW124 with *vanAB* overexpression, RW125 with combined overexpression of *vanAB* and *pcaHG*, and RW126 with stacked overexpression of *vdh*, *vanAB,* and *pcaHG* (Fig. 2B). Compared to metabolite levels in RW127 fed on FER, a decreased VAN level (by 93%, *P* < 0.001) in RW126 indicated simultaneous alleviation of the bottlenecks at both Vdh and VanAB (Fig. 2C). Compared to the wild-type strain grown on VAN, lack of PCA accumulation in RW124 and depletion of PCA (by 83%, *P* < 0.001) in RW125 demonstrated efficient PCA cleavage and thus elimination of the bottleneck at PcaHG during VAN catabolism (Fig. 2D). Cellular energy charge was 25-44% lower in the mutant strains in the coniferyl branch compared to the wild-type strain (*P* < 0.05) (Figs. 2D, 2E; SI, Table S1).

In the *p*-coumaroyl branch, in line with the reported bottleneck node at PobA^33,41,43^, there was 4HB accumulation (18.1 ± 7.6 µmol g_CDW_^-1^) during COU catabolism, while 4-hydroxybenzaldehyde (4HBA) level was 67% lower (*P* < 0.05) and PCA level was below the detection limit (Fig. 2F). Furthermore, in relation to the bottleneck, there was 31% smaller fraction of ^13^C-COU carbons in 4HB than in 4HBA within 1 min (*P* < 0.01) (Fig. 2G). During growth on 4HB, PCA accumulated (19.8 ± 5.4 µmol g_CDW_^-1^), in accordance with the reported bottleneck at PcaHG (Fig. 2F)^40^. We prepared three strains that overexpressed either *pobA* (RW128), *pcaHG* (RW123), or both *pobA* and *pcaHG* (RW129) (Fig 2H). Compared to the wild-type strain, the bottlenecks were resolved in RW128 and RW123, characterized by a 90% decrease in 4HB during growth on COU and a 50% decrease in PCA during growth on 4HB (*P* < 0.05), respectively (Figs. 2I, 2J). Similar to the coniferyl branch, the energy charge of the mutants of the *p*-coumaroyl branch was 32-40% lower than in the wild-type stain (*P* < 0.05) (Fig. 2I, 2J; SI, Table S1).

In sum, alleviation of native metabolite buildups in initial substrate catabolism led to adverse change in the cellular energy charge, thereby implying an interplay between the regulation of phenolic carbon influx and cofactor balance and a potential energy burden caused by gene overexpression^62^. Towards understanding the metabolic mechanisms underlying native cofactor balance, we investigated protein levels, metabolite pools, and the network of carbon fluxes during processing of the different phenolic compounds.

### Abundances of Transporters and Specialized Enzymes Are Modulated to Adapt to Phenolic Carbon Influx

For each of the four biological replicates, we employed a high-resolution proteomics method to capture over 4300 proteins out of the 5950 total proteins encoded by *P. putida* KT2440, including low-abundance proteins that are typically undetected (SI, Table S2). From whole-cell proteomics analysis of *P. putida* cells fed on each phenolic substrate compared to cells fed on SUC, we profiled statistically significant changes (*P* < 0.05) in the abundance of proteins involved in membrane transport, the coniferyl branch, the *p*-coumaroyl branch, and the central carbon metabolism (Fig. 3 and Fig. 4). The HcnK abundance was 700-fold greater in *P. putida* cells fed with FER or COU (Fig. 3A), in agreement with the assignment of HcnK as the common transporter for hydroxycinnamate uptake to both the coniferyl and *p*-coumaroyl branches^27^. However, HcnK abundance was also 9-fold to 13-fold higher in cells grown on VAN and 4HB relative to SUC (Fig. 3A; SI, Table S3), indicating that abundance of this transporter was also induced by downstream metabolites in the pathways. For the VAN transporter VanK^27^, its abundance was 830-fold higher in cells fed with VAN or FER compared to SUC (Fig. 3A; SI, Table S3), indicating the dependence of VanK level on the presence of VAN either as a substrate or as an intermediate in FER catabolism (Fig. 1A). Similarly, the abundance of 4HB transporter PcaK was increased to similar extent (> 140-fold) during growth across all four phenolic substrates compared to growth on SUC, in accordance with PcaK serving also as a transporter of PCA^26,27,69^, a common downstream intermediate in both the coniferyl and p-coumaroyl branches (Fig. 3A; SI, Table S3).

**Fig. 3.**
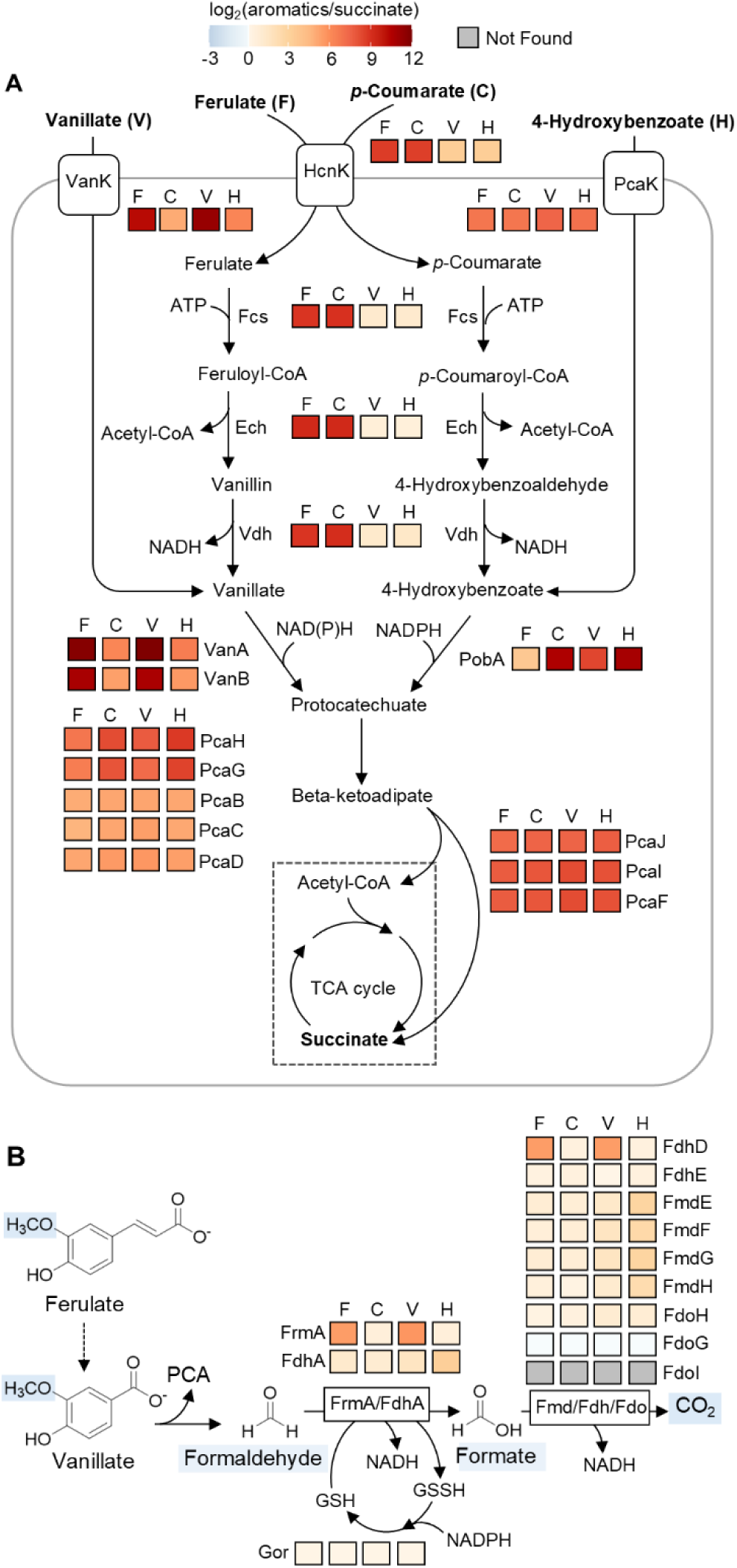
Profiling protein abundance changes in the peripheral pathways. Log_2_ fold change in protein abundance in (A) the peripheral pathways and (B) formaldehyde oxidation pathway. Abbreviation of enzymes: HcnK, hydroxycinnamic acid transporter; VanK, vanillate transporter; PcaK, 4-hydroxybenzoate/protocatechuate transporter; Fcs, feruloyl-CoA synthase; Ech, feruloyl-CoA hydratase-lyase; Vdh, vanillin dehydrogenase; VanAB, vanillate O-demethylase; PobA, 4-hydroxybenzoate 3-monooxygenase; PcaH, protocatechuate 3,4-dioxygenase beta chain; PcaG, protocatechuate 3,4-dioxygenase alpha chain; PcaC, 4-carboxymuconolactone decarboxylase; PcaB, 3-carboxy-cis,cis-muconate cycloisomerase; PcaD, 3-oxoadipate enol-lactonase; PcaIJ, 3-oxoadipate CoA-transferase; PcaF, betaketoadipyl-CoA thiolase; FrmA, formaldehyde dehydrogenase; FdhA, formaldehyde dehydrogenase; FmdEFGH, formate dehydrogenase; FdhDE, formate dehydrogenase; FdoHGI, formate dehydrogenase; Gor, glutathione reductase. Log_2_ fold change and *P* values are provided in SI, Table S3, using data obtained from four independent biological replicates (*n* = 4). The full dataset of proteomics is provided in Supplemental Dataset 1.

**Fig. 4.**
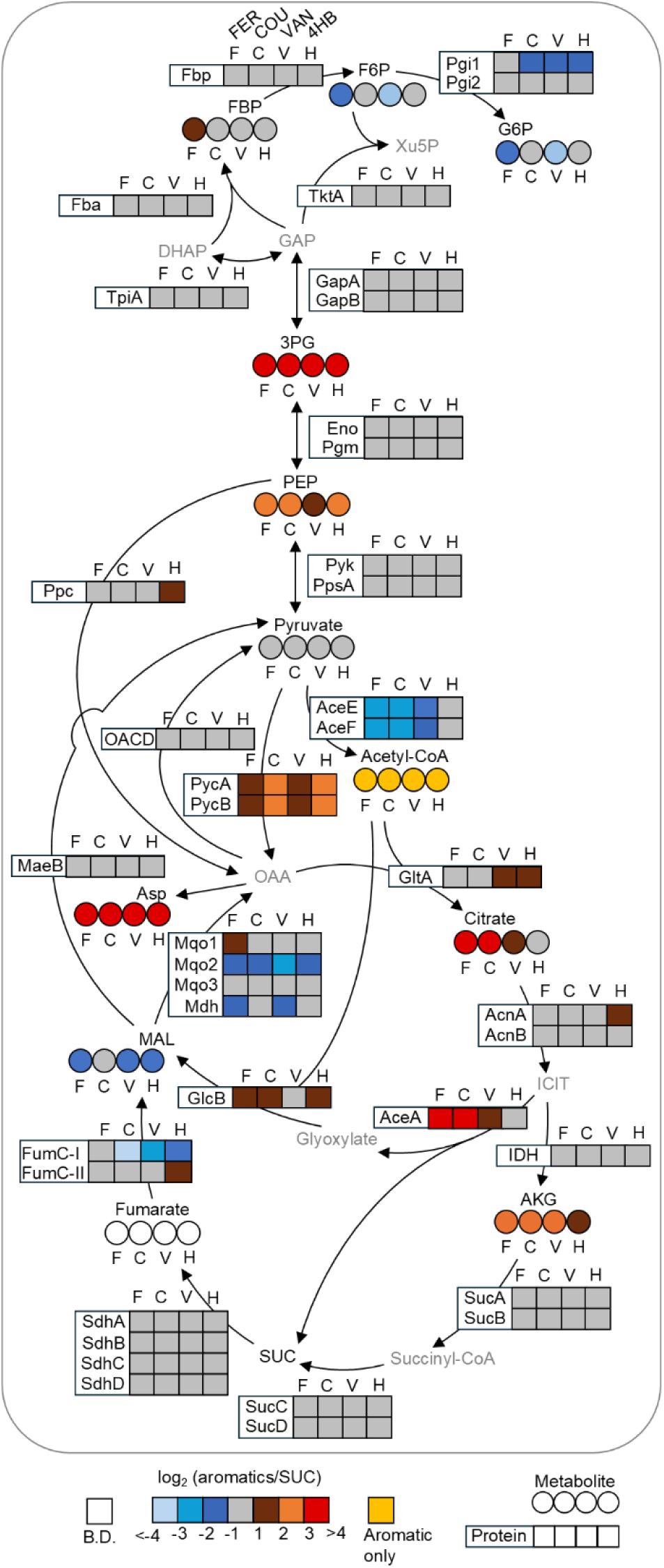
Profiling of intracellular metabolite levels and protein abundances in central carbon metabolism. Metabolite abbreviations are as mentioned in the main text. Protein abbreviations are as follows: Fbp, fructose-1,6-bisphosphatase; Pgi, glucose-6-phosphate isomerase; Fba, fructose-1,6-bisphosphate aldolase; TktA, transketolase; TpiA, triosephosphate isomerase; Gap, glyceraldehyde-3-phosphate dehydrogenase; Eno, enolase; Pgm, phosphoglucomutase; Pyk, pyruvate kinase; PpsA, phosphoenolpyruvate synthase; Ppc, phosphoenolpyruvate carboxylase; OACD, oxaloacetate decarboxylase; AceEF, pyruvate dehydrogenase; PycAB, Pyruvate carboxylase; MaeB, malic enzyme; GltA, citrate synthase; Acn, aconitate hydratase; IDH, isocitrate dehydrogenase; SucAB, oxoglutarate dehydrogenase; SucCD, succinyl-CoA ligase; FumC, fumarate hydratase; Mqo, malate:quinone oxidoreductase; Mdh, malate dehydrogenase; AceA, isocitrate lyase; GlcB, malate synthase. B.D: below detection limit. Data of the protein abundances and metabolite levels from four independent biological replicates (*n* = 4) are provided in SI, Tables S3 and S4. The full dataset of proteomics is provided in Supplemental Dataset 1.

After substrate uptake, a series of cofactor-dependent enzymes are involved in the initial catabolism (Fig. 3A). Compared to SUC-fed cells, the abundances of Fcs, Ech, and Vdh, which collectively catalyze the conversion of FER to VAN or COU to 4HB, were up to 590-fold higher in cells fed on FER or COU, but remained unchanged in cells grown on VAN or 4HB (Fig. 3A; SI, Table S3). The abundance of VanAB, which catalyze VAN *O-*demethylation to produce PCA and formaldehyde, was up to 2700-fold higher in cells grown on FER or VAN compared to SUC; the corresponding increase during feeding on COU and 4HB was 60-fold (Fig. 3A; SI, Table S3). To avoid the accumulation of toxic formaldehyde, *P. putida* can oxidize formaldehyde to CO_2_ via formate as an intermediate using formaldehyde dehydrogenase (FrmA, FdhA) and formate dehydrogenase (Fdh, Fmd, Fdo) (Fig. 3B)^70,71^. Accordingly, relative to SUC-grown cells, the abundance of FrmA and FdhD was up to 40-fold higher in cells grown on FER or VAN, but remained unchanged in cells grown on COU or 4HB (Fig. 3B). Related to stress tolerance strategies^72,73^, abundance of TtgABC efflux pump was 3-fold higher in VAN-grown cells compared to SUC-fed cells (SI, Table S3), suggesting its involvement in formaldehyde tolerance. For the conversion of 4HB to PCA, the abundance of PobA was up to 3100-fold higher during feeding on COU or 4HB relative to SUC feeding; the corresponding change was up to 550-fold higher in cells grown on FER or VAN (Fig. 3A; SI, Table S3).

Taken together, the increased abundance of the initial catabolic enzymes across all substrates implied co-regulation in the utilization of the different phenolic substrates. In fact, compared to growth on SUC, the levels of enzymes (PcaHGBCDJIF) involved in the *ortho*-cleavage of PCA via the β-ketoadipate pathway showed 40-fold to 500-fold increase during growth on the four phenolic substrates (Fig. 3A; SI, Table S3). Notably, the β-ketoadipate pathway generates SUC and acetyl-CoA as additional carbon influxes into the TCA cycle compared to metabolism of SUC alone (Fig. 3A). Next, we evaluated changes in both protein abundances and intracellular metabolite levels in the central carbon metabolism (Fig. 4).

### Fluxes through β-Ketoadipate Pathway and Enhanced Anaplerosis Promote Carbon Retention in the TCA Cycle

During phenolic substrate utilization, malate decreased by up to 70% (*P* < 0.05), while PEP and 3-phosphoglycerate (3PG) increased 2-fold to 60-fold (*P* < 0.01) relative to SUC utilization, indicating strong cataplerotic flux from the TCA cycle towards intermediates in gluconeogenesis (Fig. 4; SI, Table S4). By contrast, citrate (CIT) and alpha-ketoglutarate (AKG) levels were up to 10-fold and 7-fold higher (*P* < 0.001), respectively, in cells grown on phenolic substrates than on SUC, implying high carbon flux towards the high energy-producing side of the TCA cycle (Fig. 4; SI, Table S4). Despite these changes in metabolite levels, only three nodes in central carbon metabolism showed greater than 2-fold changes in protein abundance during growth on the phenolic substrates relative to growth on SUC. First, the enzyme catalyzes the anaplerotic reaction of pyruvate carboxylation to OAA (Pyc) was 2.5-fold to 6-fold more abundant, but the abundance of Ppc, which catalyzes the other anaplerotic reaction of PEP carboxylation remained mostly unchanged (< 2-fold), suggesting a dominant role of Pyc during growth on phenolic compounds (Fig. 4; SI, Table S3). Second, the two enzymes (AceA and GlcB) involved in the glyoxylate shunt were 2-fold to 30-fold more abundant (Fig. 4; SI, Table S3). Third, the abundance of AceE and AceF, which catalyze pyruvate to acetyl-CoA, was up to 6-fold lower in cells fed on FER or COU, potentially due to the acetyl-CoA surplus. Thus, the proteomics profiling implied an increased cataplerotic flux, activation of the glyoxylate shunt, and a decreased flux in pyruvate dehydrogenase during phenolic catabolism. To corroborate this inferred metabolic remodeling, we performed kinetic ^13^C-profiling of *P. putida* KT2440 cells switched from unlabeled substrates to 50% phenyl-^13^C_6_-FER, phenyl-^13^C_6_-COU, phenyl-^13^C_6_-VAN, phenyl-^13^C_6_-4HB, or U^13^C_4_-SUC (Fig. 5A).

**Fig. 5.**
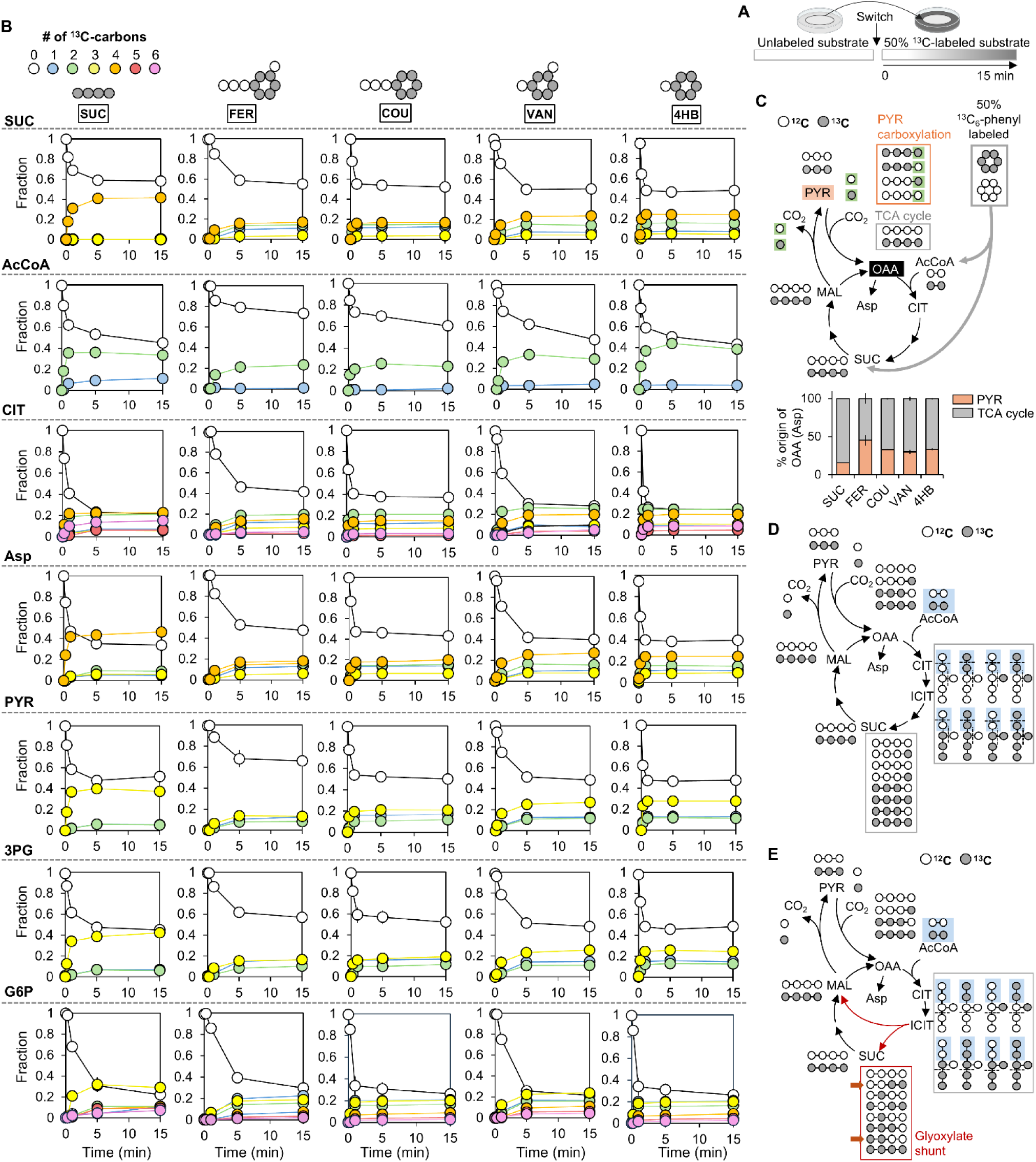
Kinetic ^13^C-profiling and carbon mapping of metabolites in the central carbon metabolism. (A) Schematic illustration of kinetic isotope incorporation experiment using 50% ^13^C-labeled substrates. (B) Kinetic ^13^C-profiling of central carbon metabolites. (C) Contribution from pyruvate versus malate to the oxaloacetate pool determined by carbon mapping when cells were grown on 50% ^13^C-labeled phenolic substrates or SUC. Isotopologues of TCA cycle intermediates predicted by carbon mapping when assuming (D) 100% canonical TCA cycle and (E) 100% glyoxylate shunt. Kinetic ^13^C-profiling of dihydroxyacetone phosphate, PEP, and sedoheptulose-7-phosphate are provided in SI, Fig. S4. For clarity, we presented the data in (B) as mean ± standard deviation from three independent biological replicates (*n* = 3), error bars are not visible where data across the three biological replicates were highly reproducible. Individual data points for kinetic ^13^C-metabolomics are provided in Supplemental Dataset 2. Abbreviations of metabolites are the same as shown in the main text.

The kinetic ^13^C-profiling of intracellular metabolites was aimed at capturing routing of substrate carbons in specific metabolic pathways. After 1 min, the ^13^C-labeled fractions of SUC were at up to 24% in cells grown on FER and VAN, but at up to 52% in cells grown on COU and 4HB (*P* < 0.001) (Fig. 5B). The slower incorporation of ^13^C into the coniferyl branch relative to the *p*-coumaroyl branch was consistent with up to 58% slower uptake rates of the respective substrates (SI, Fig. S1). The ^13^C-labeled fraction of acetyl-CoA in cells fed with the hydroxycinnamates (FER and COU) was up to 50% lower than in cells grown on the hydroxybenzoates (VAN and 4HB) (*P <* 0.05), due to the production of unlabeled acetyl-CoA via Ech (Fig. 5B). Further, consistent with the additional acetyl-CoA influx from the β-ketoadipate pathway, ^13^C-labeled acetyl-CoA was predominantly doubly ^13^C-labeled (> 90%) during growth on the phenolic substrates (Fig. 5B). Under similar uptake rates, despite 3-fold to 30-fold higher CIT and aspartate (Asp) levels in cells fed with 4HB compared to SUC (*P* < 0.001), ^13^C-labeled fraction within 15 s was up to 2-fold higher in 4HB-grown cells (*P* < 0.01), confirming exceptionally high TCA cycle flux for 4HB assimilation (SI, Table S4; Figs. 5B).

We further confirmed metabolic remodeling through carbon mapping combined with ^13^C-metabolomics (Figs. 5C, 5D, and 5E). First, regarding metabolic remodeling through anaplerosis inferred from the change of Pyc abundance, we determined the relative fraction of OAA derived from pyruvate by tracking triply ^13^C-labeled OAA (via Asp labeling), which would be generated only from the carboxylation of triply ^13^C-labeled pyruvate by unlabeled CO_2_ (Fig. 5C). At the plateau of the kinetic ^13^C data, we recorded 38% triply ^13^C-labeled pyruvate and 6% triply ^13^C-labeled Asp in the SUC-fed cells (Fig. 5B), reflecting that 16% of OAA was derived from pyruvate (Fig. 5C), consistent with previous flux analysis^74^. However, the isotopologue profiling in cells fed on the phenolic substrates revealed 30% to 45% of OAA derived from pyruvate (Fig. 5C), confirming the enhanced Pyc-catalyzed pyruvate carboxylation to OAA (*i.e.*, anaplerosis) implied by the proteomics data.

Second, to verify metabolic remodeling through the glyoxylate shunt, we illustrated carbon mapping for two scenarios. In one scenario, SUC generated exclusively through the canonical TCA cycle with no flux through the glyoxylate shunt would result in non-labeled, singly ^13^C-labeled, triply ^13^C-labeled, and fully ^13^C-labeled SUC after one cycle of carbon mapping (Fig. 5D); an additional cycle would generate the same isotopologues (SI, Fig. S3). In another scenario, the glyoxylate shunt in the TCA cycle would generate doubly ^13^C-labeled SUC through isocitrate lyase (AceA) (Fig. 5E). Detection of this doubly ^13^C-labeled isotopologue only in the cells fed on phenolic substrates confirmed the activation of the glyoxylate shunt during the metabolism of these substrates (Fig. 5B). We performed ^13^C-fluxomics to obtain quantitative insights into the metabolic network remodeling.

### High flux retention in the TCA cycle sustains cofactor surplus

The algorithm for the ^13^C-fluxomics of each of the four phenolic substrates was constrained by biomass growth rates, substrate depletion rates, metabolite secretion rates, intracellular metabolite levels, and kinetic ^13^C-profiling of metabolites (Fig. 6A-6D). Relative to SUC metabolism, the β-ketoadipate pathway provides an additional influx of acetyl-CoA from the phenolic substrates and the presence of the acryl group in the hydroxycinnamates generates a surplus acetyl-CoA (Fig. 1A). Consequently, we obtained 2-fold to 9-fold increase in carbon fluxes in the TCA cycle during phenolic catabolism compared to SUC utilization (Figs. 6A-6D; SI, Fig. S5). Moreover, the citrate synthase flux, which combines OAA with acetyl-CoA to form CIT, was up to 2-fold higher for the catabolism of the hydroxycinnamate compounds (FER and COU) than the hydroxybenzoate compounds (VAN and 4HB) (Figs. 6A-6D). Accordingly, flux through the glyoxylate shunt was determined to be 2.5-fold to 13-fold higher during metabolism of the hydroxycinnamates than the hydroxybenzoates (Figs. 6A-6D). This flux difference with the different phenolic compounds was consistent with our proteomics data on the relative abundance of AceA in the glyoxylate shunt (Fig. 4; SI, Table S3).

**Fig. 6.**
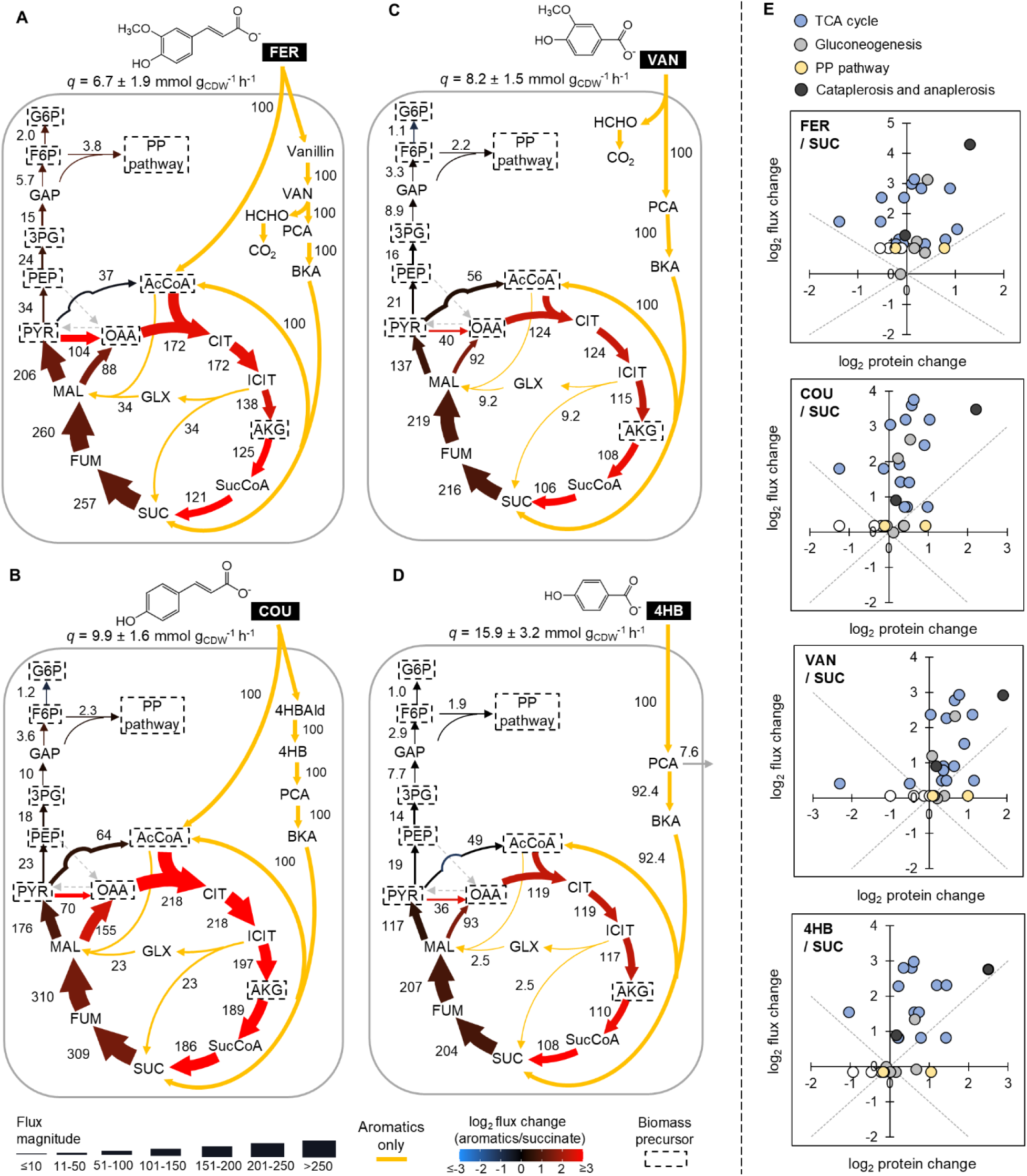
Quantitative flux analysis of phenolic acid metabolism and correlation between ^13^C-fluxomics and proteomics. Optimized flux model of *P. putida* KT2440 grown on (A) FER, (B) COU, (C) VAN, and (D) 4HB as the sole carbon source constrained by proteomics, metabolite pool sizes, and ^13^C metabolomics data. (E) Correlation matrices of relative flux changes versus relative protein abundance changes. All fluxes are normalized to the substrate uptake rate (*q*). The weight of arrows is relative to flux magnitude. Relative flux changes between cells grown on the phenolic substrates and SUC are denoted by a red-black-blue color scheme. Increased flux is shown in shades of red, decreased flux is shown in shades of blue, and black indicates negligible differences in flux values. The yellow arrows represent fluxes that are predicted to be only in phenolic acid catabolism. The grey arrow represents metabolite secretion. Flux model of *P. putida* grown on SUC is provided in SI, Fig. S5. The complete flux estimations are provided in Supplemental Dataset 3. Abbreviations of metabolites are the same as shown in the main text.

With respect to the flux remodeling at the cataplerosis-anaplerosis nodes, the cataplerotic flux through malic enzyme was 1.4-fold to 2.5-fold higher during phenolic substrate catabolism compared to SUC catabolism, while there was no flux through OAA decarboxylase (Figs. 6A-6D; SI, Fig. S5). Informed by the proteomics data, which highlighted up to 6-fold increase in Pyc (pyruvate carboxylase) relative to less than 2-fold increase in Ppc (PEP carboxylase) (*P* < 0.001), anaplerotic flux was constrained to occur only via Pyc, involving the carboxylation of pyruvate to OAA, to replenish four-carbon intermediates in the TCA cycle (Fig. 4). This pyruvate-to-OAA anaplerotic flux was 7-fold to 20-fold higher during phenolic substrate utilization than SUC utilization (Figs. 6A-6D; SI, Fig. S5), in agreement with the 2-fold to 3-fold higher OAA production from pyruvate influx illustrated by the carbon mapping combined with kinetic ^13^C-metabolomics (Fig. 5C).

For resolving whether to include flux through pyruvate dehydrogenase, we performed ^13^C-fluxomics with and without the reaction of pyruvate to acetyl-CoA (SI, Table S5). The model fit was not statistically acceptable when we deactivated the reaction in FER and COU (SI, Table S5). Therefore, pyruvate dehydrogenase was thus considered active during the phenolic substrate catabolism despite the observed decrease in the protein abundance (Fig. 4), suggesting possible sufficient abundance to support flux or post-translational regulation^18,75^.

When we compared our high-resolution ^13^C-fluxomics of the metabolic network with the widely used approach of flux balance analysis (FBA), both revealed that sustenance of high TCA cycle flux was prioritized over gluconeogenic flux during phenolic substrate catabolism (Figs. 6A-6D; Supplemental Dataset 4). Specifically, the ^13^C-fluxomics predicted that only 13-16% of the cataplerotic flux from the TCA cycle to pyruvate was invested towards gluconeogenic flux, while 69-77% was recycled back to the TCA cycle via both pyruvate carboxylase (to replenish OAA) and pyruvate dehydrogenase (to generate acetyl-CoA) (Figs. 6A-6D). The FBA still predicted 78-86% of carbon retention in the TCA cycle but indicated low cataplerotic flux through malic enzyme (14-22%) instead of the anaplerotic carbon recycling predicted by the ^13^C-fluxomics, a difference that we attributed to the low resolution of the anaplerosis-cataplerosis node in the FBA algorithm (Supplemental Dataset 4).

With respect to relationships between our ^13^C-fluxomics and proteomics data, there was a general trend of positive correlation between changes in protein abundances and changes in fluxes in cataplerosis, anaplerosis, 4 reactions in gluconeogenesis, and at least 7 reactions in the TCA cycle (Fig. 6E). Due to the discrepant magnitudes in the respective protein abundance and flux changes, there was an overall poor correlation between the relative changes in the metabolic fluxes versus changes in protein abundances (Pearson correlation, *r* = 0.11-0.35, *P* = 0.08-0.59) (Fig. 6E). Therefore, our correlation matrices are consistent with the proposal that metabolic fluxes in central carbon metabolism are driven primarily by thermodynamic factors and post-translational modifications^18,75^.

To examine consequences of the remodeled metabolic network on cofactor balance, we determined the rates of production and consumption of NADPH, NADH/UQH_2_, and ATP during the catabolism of the phenolic substrates compared to SUC, as a reference (Figs. 7A and 7B). Increased flux through cataplerosis via malic enzyme and through isocitrate dehydrogenase in the TCA cycle led to 1.2-fold to 2.5-fold greater NADPH production in cells grown on phenolic substrates compared to SUC-grown cells (Fig. 7B). As a result, the NADPH production was 24%-44% in excess of the NADPH demand for biomass synthesis and the peripheral pathway including formaldehyde detoxification; this NADPH surplus was routed to NADH synthesis through transhydrogenase reactions (Fig. 7B). Due to the promoted retention of carbon flux in the TCA cycle, NADH/UQH_2_ production was 1.4-fold to 2.7-fold higher during catabolism of phenolic substrates relative to SUC utilization (Fig. 7B). The NADH surplus was 1.4-fold to 2.7-fold higher in phenolic substrate-fed cells than SUC-grown cells, resulting in a corresponding fold increase in ATP production via oxidative phosphorylation (Fig. 7B). In corroboration of the model-predicted ATP surplus, absolute quantitation of the cellular ATP content revealed up to 2-fold higher ATP in cells fed with FER, COU, or 4HB than in the SUC-fed cells (*P* <0.01) (Fig. 7C). The surplus of ATP was consistent with the relatively higher energy charge values during processing of the phenolic substrates compared to SUC (Fig. 1C), underscoring the optimization of the carbon metabolism in *P. putida* to maintain a high-energy state during phenolic substrate utilization.

**Fig. 7.**
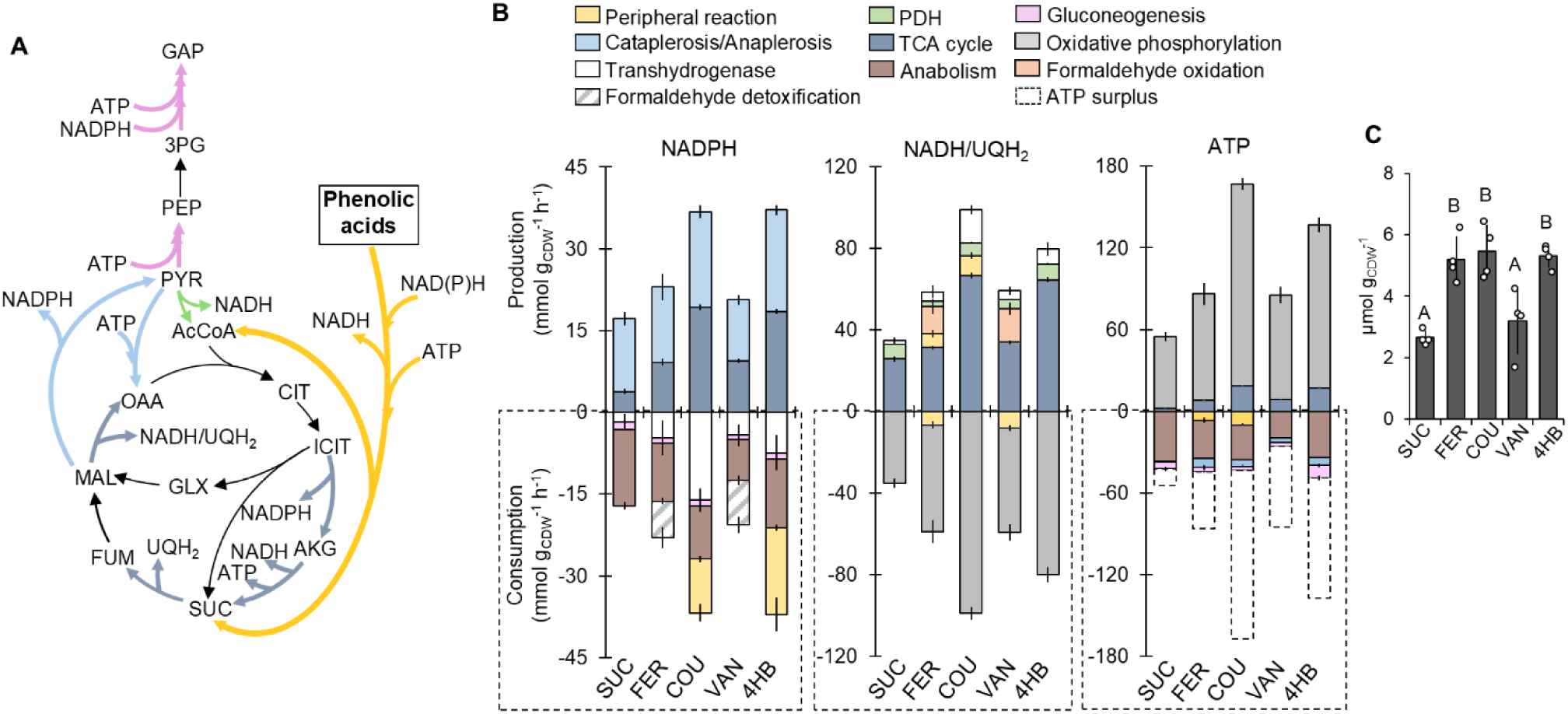
Cofactor balance in *P. putida* during carbon metabolism of the four lignin-derived phenolic substrates compared to succinate. A) Schematic illustration of metabolic reactions involving NADPH, NADH/UQH_2_, and ATP. (B) Production and consumption rates (mmol g_CDW_^-1^ h^-1^) of NADPH, NADH/UQH_2_, and ATP. Production of ATP from NADH/UQH_2_ was calculated using a phosphate to oxygen (P/O) ratio of 1.5. Abbreviations of metabolites are the same as shown in the main text. (C) Quantified intracellular ATP pool size (µmol g_CDW_^-1^) in cells grown on the lignin-related phenolic substrates compared to succinate. Data are expressed as mean ± standard deviation from four independent biological replicates (*n* = 4). In C, one-way ANOVA was performed followed by Tukey’s HSD post hoc test. Statistically significant differences (*P* < 0.05) are denoted by a change in letter.

## Discussion

The catabolism of lignin-derived phenolic substrates represents a critical step in valorizing lignin using biotechnologically relevant microbes^2–7^. Strains of *P. putida* are extensively studied for this endeavor due to their native ability to catabolize various aromatic compounds^10,11,13,14,16–18,37,76^. However, elucidation of the native metabolic network in relation to cofactor balance in *P. putida* during catabolism of phenolic substrates remains a knowledge gap. Here, we performed a comprehensive quantitative analysis of the metabolic underpinnings of carbon and energy fluxes in *P. putida* KT2440 during conversion of two hydroxycinnamates and two hydroxybenzoates, which represent common lignin-derived phenolic substrates.

To address metabolic bottlenecks during conversion of phenolic substrates, increase in enzyme expression was shown to be a successful strategy when an enzyme (i.e., VanA or VanB) exhibited a preference for NADH over NADPH, but bottleneck was not alleviated for an enzyme (i.e., PobA) with NADPH specificity due to cofactor deficiency^28,33,40,43^. The latter deficiency was circumvented by substituting for an enzyme with broader cofactor specificity (i.e., PraI) or by increasing NADPH supply through substrate co-feeding^41,43^. Our quantitative flux analysis determined that the metabolic fluxes generated excess in both NADPH and NADH in *P. putida* KT2440, but the NADH surplus was up to 6-fold higher than the NADPH surplus. Thus, our findings implied that the native cofactor balance was more amenable to buffering increased NADH demand than increased NADPH demand from enzyme overexpression.

Predicted by our proteomics data and confirmed by ^13^C-fluxomics, we identified key nodes of flux remodeling with important energetic consequences during the metabolism of the phenolic substrates. At the cataplerosis-anaplerosis nexus between the TCA cycle and lower glycolysis, *P. putida* KT2440 exhibited the specific cataplerotic routing of carbon fluxes through malic enzyme (to deplete malate and produce pyruvate) and anaplerotic routing through pyruvate carboxylase (to replenish OAA). This adaptation of cataplerosis and anaplerosis generated 50% of the NADPH yield and sustained high flux through the TCA cycle to produce over 70% of the NADH yield, which eventually fueled oxidative phosphorylation for ATP production. A similar routing of carbon fluxes through the cataplerosis-anaplerosis nodes was noted in glucose-grown *P. putida* to satisfy elevated cofactor demands in response to oxidative and energetic stresses^18,59^. For instance, in response to oxidative stress, *P. putida* has been shown to enhance carbon flux from glucose catabolism through the oxPPP to generate excess NADPH that exceeded 50% of the biomass demand^59^. Moreover, when the conversion of NADH to ATP via oxidative phosphorylation was uncoupled chemically in glucose-fed *P. putida*, there was an increase in carbon flux through the high-energy producing side of the TCA cycle towards providing NAD(P)H and ATP production to counterbalance the metabolic perturbation^54^. Thus, we propose that cofactor-driven metabolic reprogramming is a widespread phenomenon under different substrate utilization.

The *ortho*-cleavage pathway, the only known catabolic route for the phenolic substrates in *P. putida* (particularly strain KT2440), generates SUC and acetyl-CoA to feed into the TCA cycle. There are two other reported aerobic PCA cleavage pathways: for example, the 2,3-*meta* cleavage pathway found in *P. putida* strains mt-2 and H, and the 4,5-*meta* cleavage pathway revealed in *Comamonas testosteroni* and *Sphingobium sp*. SYK-6^75,77–81^. The 2,3-*meta* cleavage route generates pyruvate and acetyl-CoA with the production of NADH^78,81^; the 4,5-*meta* cleavage route generates OAA and pyruvate with the production of NADPH^75,80^. The quantitative analysis employed here to decode cofactor balance involved in the *ortho*-cleavage pathway could be used to evaluate the metabolic routing of phenolic carbons and cellular cofactor balance following the other cleavage pathways.

Here we focused on the catabolism of hydroxycinnamates and hydroxybenzoates with PCA as the common catabolic intermediate in *P. putida* KT2440. Lignin-derived compounds can constitute other aromatic structures^9^, with different intermediates such as catechol and gallate^10,13,17^. Moreover, sugar monomers and aliphatic acids often co-exist with phenolic acids in hydrolysates of lignocellulosic feedstocks^82,83^. Further investigation is needed to address the energy metabolism in *P. putida* during co-utilization of different phenolic structures with or without sugars or aliphatic acids.

In sum, we unraveled interplay between carbon metabolism and energy metabolism in *P. putida* KT2440 grown on lignin-derived phenolic substrates, thus addressing the knowledge gap of quantitative flux analysis during utilization of these substrates. Our findings provide a valuable quantitative blueprint of the native metabolic network for further explorations of Pseudomonads and other related bacterial platforms for lignin valorization.

## Materials and Methods

### Bacterial cultivation

*Pseudomonas putida* (strain KT2440) was obtained from the American Type Culture Collection (ATCC) and stored in Lysogeny broth (Miller) nutrient-rich medium with 25% glycerol at −80 °C. Cells were washed and inoculated from frozen stocks into 20-mL glass tubes that contained 5 mL of pH-adjusted (7.0) minimal nutrient medium with 100 mM C of ferulate (FER), *p*-coumarate (COU), vanillate (VAN), 4-hydroxybenzoate (4HB) or succinate (SUC) as the sole carbon source. The minimal nutrient medium contained 5.0 mM NaH_2_PO_4_, 20 mM K_2_HPO_4_, 37 mM NH_4_Cl, 17 mM NaCl, 0.81 mM MgSO_4_ · 7H_2_O, and 34 µM CaCl_2_ · 2H_2_O. Addition of essential trace metal nutrients and FeSO_4_ (30 µM) to the nutrient medium was done prior to inoculation. As previously described ^60^, after initial inoculation of the culture in glass tubes at 30 °C with shaking at 220 rpm overnight in an incubator-shaker (New Brunswick Scientific, Edison, NJ). The cells were harvested, washed and subsequently transferred into 125-mL or 250-mL baffled flasks. To optimize oxygen transfer, only one-fifth of the volume of the baffled flask was filled with the minimal nutrient medium with carbon source. Cell growth was monitored by measuring the optical density at 600 nm (OD_600_) using an Agilent Cary UV-Vis spectrophotometer (Santa Clara, CA). Cell dry weight per gram (g_CDW_) was measured by weighing the mass of lyophilized cell pellets collected throughout the growth as previously described^60^. The conversion factor of OD_600_ and g_CDW_ was calculated through linear regression with *R*^2^ coefficient greater than 0.80 as previously described^60,84^. All growth experiments were conducted in three biological replicates.

### Mutant construction

Construction details for strains, oligonucleotides, and plasmids are detailed in Tables S6-S8. In brief, all plasmids were synthesized and cloned by Twist Biosciences. Ribosome binding sites were optimized for each native *P. putida* KT2440 sequence as described previously^85^. For the construction of *P. putida* strains, the parent strain *P. putida* AG5577 containing a total of nine unique attB sites was used as described previously^86^. The parental strain *P. putida* AG5577 was confirmed to be identical to the wild-type *P. putida* KT2440 in phenolic acid utilization (Fig. S2). All integrations were achieved by serine recombinase-assisted genome engineering (SAGE) using either the BxB1 or TG1 recombinase as previously established^86–88^. Integrations were confirmed using Oxford Nanopore sequencing at Plasmidsaurus (https://www.plasmidsaurus.com).

### Measuring extracellular substrate depletion

Aliquots of cell suspensions were collected and spanned at different time points throughout biomass growth. Following filtering during centrifugation (Costar Spin-X, 0.22-µm-pore-size filter), the filtrates were stored at −20 °C until analysis. The concentration of each aromatic substrate was determined using ultra-high-performance liquid chromatography (UHPLC) with an Agilent ZORBAX Eclipse Plus C18 column (4.6 × 100 mm with 5 µm particle size) and a UV detector at 210 nm for 4HB and VAN detection, and 275 nm for FER and COU detection. The injection volume was 10 µL and the column was maintained at 25 °C. The mobile phase consisted of 0.1% formic acid (eluent A) and 80:10:10 acetonitrile: methanol: H_2_O (eluent B) with a flow rate of 0.9 mL · min^-1^. The gradient of eluent B was set as 0 min, 6%; 6.5 min, 15%; 8.5 min, 25%; 10.75 min, 37.5%; 15.5 min, 65%; 16.25 min, 6%. Extracellular substrate depletion rate, in mmol g_CDW_^-1^ h^-1^, was determined by regression analysis. The rate of SUC depletion was obtained by Wilkes et al.^60^

### Quantitative Proteomics

*Pseudomonas putida* KT2440 was cultured until mid-exponential phase (OD_600_ = 1.0-1.2) using 100 mM C of FER, COU, VAN, 4HB or SUC as the sole carbon source in the nutrient minimal medium. After centrifugation of 2-mL cultures (10000 *g* for 5 min at 4 °C), the cell pellets were washed twice with phosphate-buffered solution (pH = 7.4) followed by quenching with cold methanol (4 °C) and incubated on ice for 30 min. The quenched cell suspensions were centrifuged, and the pellets were resuspended in methanol and stored at −80 °C until analysis.

The samples were processed with an on-filter in-cell (OFIC) digestion approach as described previously^89^. Briefly, after transferring the samples to E3 filters (CDS Analytical, Oxford, PA), and centrifuging (400 *g* for 1 min), the filter was washed once with methanol, and treated with 10 mM Tris(2-carboxyethyl)phosphineand and 40 mM chloroacetamide (in 50mM triethylammonium bicarbonate, TEAB), followed by incubating at 45 °C for 10 min. The sample-containing filters were spun to remove liquid, followed by washing once with 50mM TEAB. Subsequently, the samples were digested with Trypsin/LysC mix at 37 °C for 16-18 h. After digestion, the peptides were eluted, pooled, and dried in SpeedVac. The peptides were desalted with StageTips (CDS Analytical, Oxford, PA), dried, and stored at −80 °C until further analysis.

The peptides were first loaded onto a trap column (PepMap100 C18, 300 μm × 2 mm with 5 μm particle size; Thermo Scientific) then separated through an analytical column (PepMap100 C18, 50 cm × 75 μm with 3 μm particle size; Thermo Scientific) with eluents consisting of 0.1% formic acid in water (eluent A) and 0.1% formic acid in acetonitrile (eluent B) with a flow rate of 250 nL · min^-1^ using an Ultimate 3000 RSLCnano system with nano electrospray ionization for liquid chromatography mass spectrometry analysis (LC-MS). The MS data were acquired on an Orbitrap Eclipse mass spectrometer with FAIMS Pro Interface (Thermo Scientific) at 120K resolution, followed by MS/MS acquisition in data-independent mode following a protocol described previously^90^.

The mass spectrometry data were processed using Spectronaut software (version 19.1)^91^ and a library-free DIA analysis workflow with directDIA+ and the *P. putida* KT2440 protein sequence (Uniprot 2024 release; 5950 sequences). In short, the settings for Pulsar and library generation included: Trypsin/P as specific enzyme; peptide length from 7 to 52 amino acids; allowing 2 missed cleavages; toggle N-terminal M turned on; Carbamidomethyl on C as fixed modification; Oxidation on M and Acetyl at protein N-terminus as variable modifications; FDRs at PSM, peptide and protein level all set to 0.01; Quantity MS level set to MS2, and cross-run normalization turned on. Bioinformatics analyses were performed using Perseus software (version 1.6.2.3)^92^. The MS raw files obtained for this study have been deposited to the MassIVE server (https://massive.ucsd.edu/) with the accession number MSV000095008. The full dataset of proteomics is provided in Supplemental Dataset 1.

### Intracellular metabolite levels and isotope labeling kinetics

To profile the intracellular metabolites of each of the substrate condition (100 mM C of FER, COU, VAN, 4HB or SUC), cell suspensions (3 mL) of *P. putida* KT2440 during mid-exponential growth phase (OD_600_ = 1.0-1.2) were filtered (0.22 µm nylon membranes; Whatman, 7402-004), then lysed in a cold quenching solution (4 °C) containing methanol, acetonitrile, and water in a 2:2:1 ratio. The suspensions were centrifuged (10000 × *g*, 4 °C for 5 min) and aliquots (200 µL) of the supernatants in 2mL-tubes were dried under ultrapure nitrogen gas followed by storage at −80 °C before further analysis. The dried samples, collected from four biological replicates, were resuspended in 100 µL LC-MS grade water before metabolomics analysis.

In preparation for kinetic ^13^C-labeling experiments, 3-mL exponentially-growing *P. putida* KT2440 cells (two transfers, three biological replicates) on each unlabeled substrate (100 mM C of FER, COU, VAN, 4HB, or SUC) were filtered (0.22 µm nylon membranes; Whatman, 7402-004), and the filters containing the cells were placed onto agar plates containing 100 mM C of the same substrate, followed by incubation at 30 °C until exponential growth was resumed. To initiate ^13^C-labeling of intracellular metabolites, cell-containing filters were transferred onto fresh agar plates with 50% of ^13^C-labeled substrate: ^13^C_6_-phenyl-FER, ^13^C_6_-phenyl-COU, ^13^C_6_-phenyl-VAN, ^13^C_6_-phenyl-4HB, or U^13^C_4_-succinate. These labeled substrates were purchased from Cambridge Isotopes (Tewksbury, MA) or Sigma-Aldrich (St. Louis, MO). At four different time points (15 s, 1 min, 5 min, and 15 min), cells adhered to the filter were quenched into the aforementioned cold quenching solution (4 °C); data for time 0 were obtained with cells directly quenched from the unlabeled plate. The quenched cell suspensions were processed following the aforementioned nitrogen drying procedure and the extracted metabolites were stored at −80 °C until metabolomics analysis.

### Metabolomics analysis

Intracellular and extracellular metabolites were quantified based on our established metabolomics protocol using UHPLC (Thermo Fisher Scientific Dionex UltiMate 3000) coupled with high-resolution mass spectrometry (Thermo Fisher Scientific Q Exactive quadrupole-Orbitrap) operating in negative mode^60,93^. The LC column, an Acquity UPLC BEH C18 Column (particle size 1.7 µm, 2.1 mm × 100 mm, Waters), was maintained at 25 °C with an injection volume of 10 µL. The eluents were adopted as previously described with a flow rate of 0.18 mL·min^-193^. The data was analyzed on the Thermo Scientific XCalibur software (v3.0) using a series of standard concentrations. The fraction of different isotopomers was analyzed using the Metabolomic Analysis and Visualization Engine (MAVEN) software version 2011.6.17^94^; to correct for the natural abundance of ^13^C, IsoCor v2.2.0^95^ was used. The MS data for metabolomics have been deposited to MetaboLights depository (https://www.ebi.ac.uk/metabolights/) under the accession number of MTBLS11484. The full dataset of kinetic ^13^C-metabolomics is provided in Supplemental Dataset 2.

### 13C-fluxomics analysis

We employed both the levels and the kinetic isotopic profiling of intracellular metabolites to perform an isotope non-steady state metabolic flux analysis (INST-MFA) ^96^. A core *P. putida* metabolic network was constructed based on published resources with modifications to supplement aromatics utilization pathways^58,61^. The biomass yield equation was modified from a previous genome scale model for *P. putida* KT2440^97^. The INCA software (v2.3) based on Matlab platform^98^ was used to simulate isopotomer balances when *P. putida* was fed on different aromatic substrates and succinate. The INST-MFA models were constrained with growth, substrate consumption and metabolite secretion rates, measured pool sizes (CIT, AKG, SUC, PYR, PEP, 3PG, G6P, AcCoA) and kinetic isotopomer distributions (CIT, AKG, SUC, OAA as Asp, PEP, PYR, 3PG, DHAP, G6P, S7P, AcCoA). Flux estimations were reiterated at least 50 times from random initial values as previously suggested to obtain the best fits^99^. At least three independent flux estimations were performed for each INST-MFA model to ensure the flux estimations were consistent. The estimates were considered as statistically acceptable fits when the results passed the χ^2^ goodness-of-fit test (cutoff 95% confidence level), i.e., the minimized variance-weighted sum of squared residuals (SSR) were within the expected range. The standard errors (95% confidence intervals) of the flux estimations for the reactions in the central carbon metabolism were calculated through Monte Carlo simulations as previously reported^61,98^. Complete flux estimations are in Supplemental Dataset 3. Cofactor balance was calculated from quantified fluxes for metabolic reactions with production or consumption of NADPH, NADH/UQH_2_, and ATP. Only NADH was included for vanillate *O*-demethylase because VanB was shown to prefer NADH over NADPH^28^.

### Flux balance analysis

The most recent genome-scale model of *P. putida* KT2440 (iJN1463) was downloaded from the BIGG database (http://bigg.ucsd.edu/)^97^. The FER, VAN, and 4HB catabolic reactions are present in the original iJN1463 model. The COU uptake and initial catabolic reactions to coumaroyl-CoA were added manually to the model. The optimized flux distribution predicted by flux balance analysis was performed using the COBRApy library (version 0.29.1)^100^. Substrate uptake rates for SUC, FER, COU, VAN, and 4HB were fixed at the experimentally determined values (in mmol/g_CDW_/h), which are 16.0, 6.7, 9.9, 8.2, and 15.9, respectively. For the 4HB analysis, the PCA secretion rate was fixed at 1.2 mmol/g_CDW_/h as experimentally measured. Results for flux balance analysis are in Supplemental Dataset 4.

### Statistical analysis

Unpaired two-tailed *t*-test was applied to evaluate the significance of differences between two groups. One-way analysis of variance (ANOVA) was used to evaluate differences among three or more conditions. Tukey’s honestly significant difference (HSD) post hoc test was performed for pairwise comparison in addition to one-way ANOVA. The *P*-value threshold for statistically significant difference was set at 0.05.

## Supporting information

Supplemental Infomation

## Acknowledgments

This material is based upon work supported by the U.S. Department of Energy (DOE), Office of Science, Office of Biological and Environmental Research, Genomic Science Program under Award Number DE-SC0022181. This work was authored in part by the National Renewable Energy Laboratory, operated by Alliance for Sustainable Energy, LLC, for the U.S. DOE under Contract No. DE-AC36-08GO28308. We thank Jeffrey Czajka at Pacific Northwest National Laboratory and Shawn Xiao of Dr Yinjie Tang’s lab at Washington University in St. Louis for helpful advice during ^13^C-fluxomics modeling using the INCA software. We are grateful to Dr Pablo I. Nikel of the Technical University of Denmark (DTU) for helpful discussions. The views expressed in the article do not necessarily represent the views of the DOE or the U.S. Government. The U.S. Government retains, and the publisher, by accepting the article for publication, acknowledges that the U.S. Government retains a nonexclusive, paid-up, irrevocable, worldwide license to publish or reproduce the published form of this work, or allow others to do so, for U.S. Government purposes.

## Author contributions

N.Z., R.A.W., X.C., K.P.T., J.A.B., and Y.Y. performed research. G.T.B and A.Z.W provided resources. N.Z. and L.A. designed research, analyzed data, and wrote manuscript. L.A. supervised research.

## Data and materials availability

Proteomics data and metabolomics MS data are freely available in the MassIVE server (https://massive.ucsd.edu/) and MetaboLights depository (https://www.ebi.ac.uk/metabolights/) under accession numbers MSV000095008 and MTBLS11484, respectively.

## Supplemental Materials

The SI document contains:

Figures S1-S5

Tables S1-S8

Supplemental Dataset 1

Supplemental Dataset 2

Supplemental Dataset 3

Supplemental Dataset 4

