## Supplemental Infomation for "Quantitative Analysis of Coupled Carbon and Energy Metabolism for Lignin Carbon Utilization in *Pseudomonas putida*"

*[1]Department of Civil and Environmental Engineering, McCormick School of Engineering and Applied Science, Northwestern University, Evanston, IL 60208, USA; [2]Renewable Resources and Enabling Sciences Center, National Renewable Energy Laboratory, Golden, CO, USA; [3]Northwestern Center for Synthetic Biology, Northwestern University, Evanston, IL 60208, USA;[4]Department of Chemistry and Biochemistry, University of Delaware, Newark, DE 19716, USA*

This document contains:

Supplementary Figures: 5

Supplementary Tables: 8

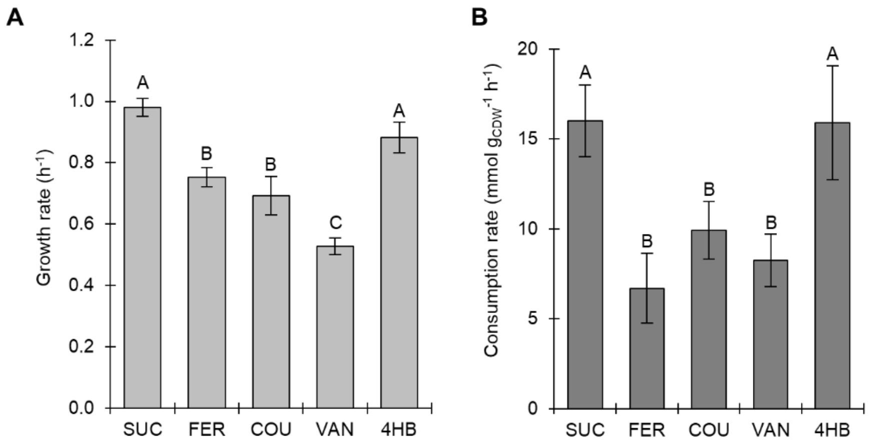


**Fig. S1**. Growth rate (A) and substrate consumption rate (B) of *P. putida* KT2440 fed with 100 mM C of SUC, FER, COU, VAN, or 4HB as the sole carbon source. In A and B, one-way analysis of variance (ANOVA) was performed followed by Tukey's HSD post hoc test. Statistically significant differences ($P < 0.05$) are denoted by a change in letter.

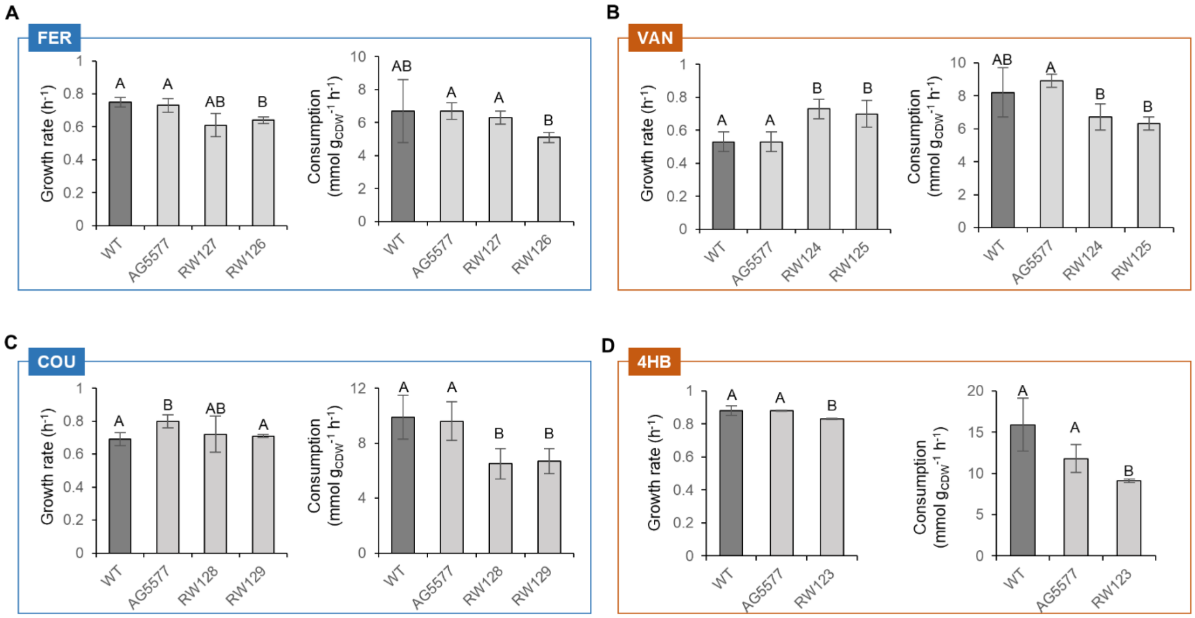


**Fig. S2.** Growth and substrate consumption rates of engineered strains on (A) FER, (B) VAN, (C) COU, and (D) 4HB. In A, B, C, and D, one-way ANOVA was performed followed by Tukey's HSD post hoc test. Statistically significant differences ($P < 0.05$) are denoted by a change in letter.

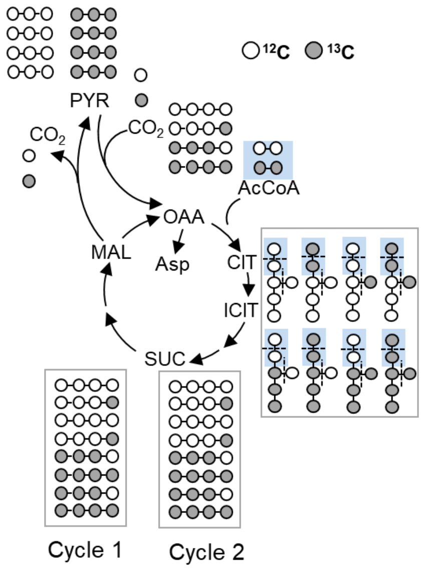


**Fig. S3.** Additional cycle of carbon mapping of the canonical TCA cycle.

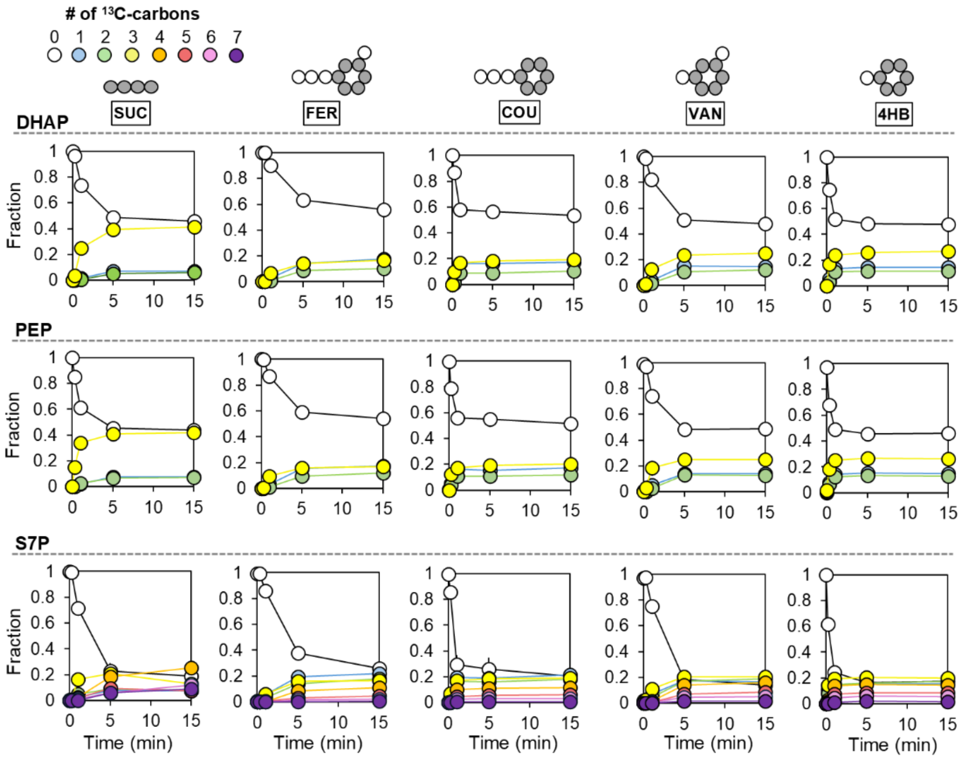


**Fig. S4.** Kinetic [13]C-profiling of dihydroxyacetone phosphate, PEP, and sedoheptulose-7-phosphate.

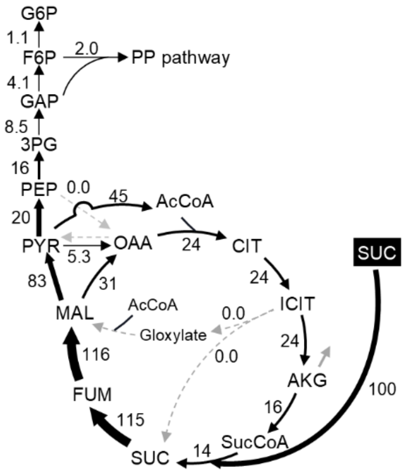


**Fig. S5**. Optimized flux model of *P. putida* KT2440 grown on SUC as the sole carbon source. The grey solid arrow represents metabolite secretion.

**Table S1**. Quantified intracellular levels (μmol $g_{CDW}^{-1}$) of ATP, ADP, and AMP and calculated energy charge values when *P. putida* cells were fed with different phenolic acid structures.

| | | FER | COU | VAN | 4HB |
|---|---|---|---|---|---|
| Wild-type strain | ATP | 5.19 ± 0.76 | 5.45 ± 0.87 | 3.69 ± 0.48 | 5.31 ± 0.37 |
| | ADP | 1.13 ± 0.23 | 0.85 ± 0.12 | 0.42 ± 0.12 | 0.74 ± 0.13 |
| | AMP | 0.46 ± 0.07 | 0.40 ± 0.01 | 0.20 ± 0.02 | 0.26 ± 0.04 |
| | Energy charge | 0.85 ± 0.01 | 0.88 ± 0.01 | 0.89 ± 0.02 | 0.90 ± 0.01 |
| AG5577 (parental strain) | ATP | 1.49 ± 0.06 | 2.16 ± 0.50 | 1.08 ± 0.10 | 3.47 ± 0.78 |
| | ADP | 0.18 ± 0.06 | 0.58 ± 0.10 | 0.12 ± 0.03 | 0.81 ± 0.09 |
| | AMP | 0.68 ± 0.13 | 0.44 ± 0.10 | 0.25 ± 0.07 | 0.91 ± 0.15 |
| | Energy charge | 0.68 ± 0.05 | 0.77 ± 0.02 | 0.79 ± 0.03 | 0.74 ± 0.05 |
| RW127 ($P_{tac}$:*vdh*) | ATP | 2.05 ± 0.32 | -- | -- | -- |
| | ADP | 0.59 ± 0.01 | -- | -- | -- |
| | AMP | 2.18 ± 0.39 | -- | -- | -- |
| | Energy charge | 0.49 ± 0.04 | -- | -- | -- |
| RW126 ($P_{tac}$:vdh: *vanAB*:*pcaHG*) | ATP | 1.12 ± 0.21 | -- | -- | -- |
| | ADP | 0.23 ± 0.06 | -- | -- | -- |
| | AMP | 1.21 ± 0.24 | -- | -- | -- |
| | Energy charge | 0.48 ± 0.07 | -- | -- | -- |
| RW124 ($P_{tac}$:*vanAB*) | ATP | -- | -- | 1.69 ± 0.48 | -- |
| | ADP | -- | -- | 0.20 ± 0.03 | -- |
| | AMP | -- | -- | 1.10 ± 0.32 | -- |
| | Energy charge | -- | -- | 0.60 ± 0.05 | -- |
| RW125 ($P_{tac}$:*vanAB* :*pcaHG*) | ATP | -- | -- | 1.28 ± 0.40 | -- |
| | ADP | -- | -- | 0.19 ± 0.04 | -- |
| | AMP | -- | -- | 0.56 ± 0.23 | -- |
| | Energy charge | -- | -- | 0.67 ± 0.12 | -- |
| RW128 ($P_{tac}$:*pobA*) | ATP | -- | 0.88 ± 0.21 | -- | -- |
| | ADP | -- | 0.30 ± 0.04 | -- | -- |
| | AMP | -- | 0.44 ± 0.07 | -- | -- |
| | Energy charge | -- | 0.63 ± 0.02 | -- | -- |
| RW129 ($P_{tac}$:*pobA*:*pcaHG*) | ATP | -- | 0.68 ± 0.07 | -- | -- |
| | ADP | -- | 0.29 ± 0.02 | -- | -- |
| | AMP | -- | 0.42 ± 0.08 | -- | -- |
| | Energy charge | -- | 0.59 ± 0.04 | -- | -- |
| RW123 ($P_{tac}$:*pcaHG*) | ATP | -- | -- | -- | 0.66 ± 0.07 |
| | ADP | -- | -- | -- | 0.31 ± 0.03 |
| | AMP | -- | -- | -- | 0.53 ± 0.07 |
| | Energy charge | -- | -- | -- | 0.54 ± 0.02 |

Note: The intracellular metabolite levels were obtained from four biological replicates.

**Table S2**. Summary of proteomics

| Condition | Replicate | # of identified proteins |
|---|---|---|
| FER | 1 | 4357 |
| | 2 | 4356 |
| | 3 | 4390 |
| | 4 | 4377 |
| COU | 1 | 4332 |
| | 2 | 4319 |
| | 3 | 4342 |
| | 4 | 4363 |
| VAN | 1 | 4383 |
| | 2 | 4363 |
| | 3 | 4357 |
| | 4 | 4349 |
| 4HB | 1 | 4355 |
| | 2 | 4361 |
| | 3 | 4364 |
| | 4 | 4357 |
| SUC | 1 | 4335 |
| | 2 | 4312 |
| | 3 | 4326 |
| | 4 | 4311 |

**Table S3**. Fold changes (log2 transformed) in protein abundances and associated *P* values in aromatic-grown relative to succinate-grown *P. putida* KT2440.

| Protein name | FER/SUC | | COU/SUC | | VAN/SUC | | 4HB/SUC | |
|---|---|---|---|---|---|---|---|---|
| | $Log_2FC$ | *P* value | $Log_2FC$ | *P* value | $Log_2FC$ | *P* value | $Log_2FC$ | *P* value |
| HcnK | 9.6 | 7.9E-12 | 9.5 | 1.1E-11 | 3.7 | 2.7E-14 | 3.1 | 1.8E-25 |
| Fcs | 9.2 | 1.3E-08 | 9.2 | 2.1E-11 | 1.2 | 5.5E-34 | 1.0 | 3.2E-31 |
| Ech | 9.0 | 3.3E-11 | 9.2 | 1.7E-10 | 0.6 | 2.9E-36 | 0.5 | 1.7E-22 |
| Vdh | 8.7 | 2.4E-07 | 8.8 | 4.1E-07 | 1.1 | 5.3E-25 | 1.3 | 4.2E-27 |
| VanK | 9.7 | 1.9E-12 | 4.2 | 1.5E-18 | 10.6 | 1.9E-08 | 5.4 | 9.4E-10 |
| VanA | 10.9 | 3.1E-10 | 5.3 | 1.2E-16 | 11.4 | 1.1E-09 | 5.5 | 4.3E-19 |
| VanB | 11.3 | 8.3E-13 | 6.0 | 3.9E-17 | 11.2 | 2.4E-09 | 6.0 | 2.7E-18 |
| PcaK | 7.1 | 1.6E-11 | 7.3 | 3.2E-13 | 7.8 | 7.7E-12 | 7.3 | 2.2E-12 |
| PobA | 4.4 | 7.2E-21 | 11.4 | 7.8E-13 | 9.1 | 2.0E-18 | 11.6 | 1.0E-12 |
| PcaH | 7.0 | 1.3E-12 | 8.5 | 4.0E-10 | 7.9 | 8.2E-16 | 9.0 | 9.3E-12 |
| PcaG | 6.5 | 4.7E-12 | 8.1 | 1.3E-09 | 7.4 | 1.6E-11 | 8.6 | 8.9E-11 |
| PcaB | 5.2 | 6.6E-14 | 5.3 | 1.6E-10 | 5.4 | 3.1E-12 | 5.3 | 1.2E-13 |
| PcaC | 5.0 | 1.5E-09 | 5.5 | 3.2E-11 | 5.7 | 2.1E-09 | 5.4 | 4.2E-12 |
| PcaD | 5.0 | 1.3E-13 | 5.4 | 7.5E-13 | 5.5 | 3.2E-13 | 5.3 | 6.5E-15 |
| PcaJ | 7.2 | 9.4E-14 | 7.5 | 1.2E-10 | 7.4 | 1.0E-11 | 7.5 | 8.4E-12 |
| PcaI | 7.4 | 6.4E-13 | 7.5 | 9.6E-13 | 7.7 | 1.9E-09 | 7.6 | 4.7E-12 |
| PcaF | 7.6 | 8.2E-11 | 7.8 | 1.1E-09 | 8.2 | 4.8E-14 | 7.9 | 7.0E-09 |
| SdhA | -0.2 | 3.7E-04 | 0.3 | 1.4E-04 | 0.3 | 1.7E-05 | 0.1 | 6.8E-03 |
| SdhB | -0.2 | 1.5E-03 | 0.3 | 2.3E-06 | 0.2 | 1.2E-04 | 0.2 | 1.8E-04 |
| SdhC | -0.4 | 6.8E-04 | 0.1 | 3.1E-01 | 0.1 | 1.5E-01 | 0.0 | 4.4E-01 |
| SdhD | 0.1 | 7.8E-01 | 0.6 | 2.2E-05 | 0.5 | 4.9E-05 | 0.3 | 3.0E-03 |
| SdhE | -0.2 | 1.6E-02 | 0.1 | 3.0E-01 | 0.4 | 4.4E-03 | -0.1 | 5.9E-03 |
| FumC-II | 0.7 | 1.2E-09 | 0.5 | 3.6E-06 | 0.6 | 9.4E-09 | 1.5 | 2.6E-05 |
| FumC-I | 0.6 | 6.1E-04 | -3.1 | 4.4E-07 | -2.6 | 1.1E-06 | -1.3 | 6.9E-06 |
| Mqo1 | 1.0 | 2.5E-11 | 0.9 | 2.5E-06 | 0.8 | 1.6E-11 | 0.5 | 3.8E-08 |
| Mqo2 | -1.7 | 3.3E-12 | -1.5 | 3.1E-06 | -2.8 | 3.6E-09 | -1.3 | 1.1E-10 |
| Mqo3 | -0.4 | 4.9E-04 | -0.1 | 8.4E-01 | -0.4 | 7.5E-04 | 0.8 | 1.4E-04 |
| Mdh | -1.4 | 1.9E-06 | 0.0 | 2.7E-02 | -1.0 | 2.5E-06 | 0.7 | 8.2E-03 |
| GltA | 0.7 | 7.9E-11 | 0.8 | 2.0E-10 | 1.0 | 9.7E-13 | 1.0 | 1.3E-13 |
| AcnA-I | 0.3 | 1.4E-06 | 0.3 | 5.6E-04 | -0.1 | 3.8E-01 | 1.5 | 1.2E-04 |
| AcnA-II | 0.0 | 3.5E-02 | 0.3 | 2.3E-04 | 0.1 | 2.5E-01 | 0.5 | 5.4E-08 |
| Idh | -0.6 | 2.0E-08 | -0.1 | 1.5E-02 | 0.4 | 1.4E-06 | 0.1 | 8.3E-03 |
| SucA | 0.0 | 6.8E-01 | 0.5 | 2.3E-09 | 0.6 | 3.3E-09 | 0.5 | 2.5E-09 |
| SucB | -0.1 | 8.1E-02 | 0.3 | 3.1E-05 | 0.4 | 5.4E-07 | 0.3 | 5.8E-06 |
| SucC | 0.1 | 1.5E-01 | 0.6 | 2.5E-06 | 0.8 | 7.1E-08 | 0.5 | 4.2E-05 |
| SucD | 0.3 | 9.6E-06 | 0.8 | 1.2E-07 | 0.9 | 6.0E-11 | 0.8 | 2.3E-07 |
| AceA | 4.9 | 2.3E-08 | 3.9 | 4.1E-08 | 1.4 | 7.4E-21 | 1.0 | 3.2E-14 |
| GlcB | 1.9 | 1.6E-09 | 1.9 | 1.9E-08 | 0.8 | 4.8E-13 | 1.2 | 3.9E-14 |
| MaeB | -0.2 | 1.9E-07 | 0.0 | 1.7E-03 | 0.2 | 9.2E-05 | 0.0 | 1.2E-03 |
| PycB | 1.3 | 2.7E-15 | 2.1 | 1.6E-13 | 1.8 | 6.6E-11 | 2.5 | 1.9E-11 |
| PycA | 1.4 | 2.7E-13 | 2.2 | 1.4E-15 | 2.0 | 1.2E-08 | 2.5 | 1.7E-09 |
| AceE | -2.5 | 5.4E-08 | -2.3 | 1.1E-08 | -1.0 | 8.6E-06 | -0.7 | 5.9E-05 |
| AceF | -2.7 | 5.6E-07 | -2.6 | 3.7E-07 | -1.4 | 1.8E-05 | -1.0 | 8.2E-05 |
| Ppc | 0.9 | 1.1E-10 | 0.8 | 7.7E-05 | 0.7 | 8.2E-09 | 1.3 | 2.6E-12 |
| Eno | -0.2 | 1.7E-03 | 0.1 | 4.1E-01 | 0.2 | 9.4E-03 | -0.2 | 1.5E-03 |
| Pgm | 0.0 | 3.4E-01 | -0.1 | 1.1E-03 | -0.3 | 8.0E-07 | -0.1 | 4.5E-05 |
| Pgk | -0.3 | 2.3E-05 | -0.3 | 9.7E-06 | -0.3 | 2.2E-05 | -0.4 | 7.1E-06 |
| GapA | 0.0 | 1.2E-03 | 0.3 | 5.3E-04 | -0.6 | 1.0E-11 | 0.9 | 1.5E-06 |
| GapB | 0.1 | 3.6E-02 | 0.3 | 6.2E-05 | 0.1 | 6.4E-01 | 0.0 | 1.9E-03 |
| TpiA | -0.3 | 2.2E-08 | -0.4 | 2.0E-08 | -0.3 | 5.0E-10 | -0.4 | 5.4E-11 |
| Fba | -0.2 | 3.6E-04 | -0.1 | 1.8E-03 | -0.1 | 1.3E-02 | -0.3 | 1.6E-06 |

| | | | | | | | | |
|---|---|---|---|---|---|---|---|---|
| Fbp | -0.5 | 9.3E-05 | -0.5 | 2.5E-05 | -0.5 | 3.1E-05 | -0.6 | 9.1E-06 |
| Pgi1 | -0.7 | 4.0E-04 | -1.1 | 1.8E-04 | -1.3 | 5.9E-05 | -1.1 | 1.3E-04 |
| Pgi2 | 0.2 | 1.0E-03 | 0.1 | 3.6E-01 | 0.0 | 4.3E-01 | 0.3 | 4.0E-04 |
| TktA | -0.4 | 1.3E-05 | -0.3 | 2.0E-05 | -0.1 | 4.3E-03 | -0.4 | 2.0E-05 |
| OACD | 0.1 | 3.4E-03 | -0.1 | 4.0E-02 | 0.8 | 1.7E-08 | -0.4 | 9.1E-06 |
| Pyk | 0.3 | 1.4E-04 | 0.2 | 2.9E-04 | 0.2 | 1.6E-02 | 0.5 | 6.2E-06 |
| PpsA | 0.3 | 3.8E-09 | 0.4 | 1.4E-09 | 0.5 | 4.6E-11 | 0.5 | 1.1E-10 |
| FrmA | 5.1 | 1.9E-10 | 0.7 | 1.8E-23 | 5.3 | 1.2E-11 | 0.6 | 3.1E-19 |
| FdhA | 1.3 | 1.3E-08 | 1.1 | 1.7E-06 | 1.7 | 6.5E-15 | 3.3 | 5.9E-07 |
| FdhD | 4.6 | 7.2E-11 | 0.3 | 1.5E-13 | 4.8 | 4.3E-10 | -0.1 | 7.3E-10 |
| FdhE | 0.5 | 1.2E-03 | 0.6 | 1.2E-03 | 0.2 | 4.3E-02 | 0.6 | 1.3E-03 |
| FmdE | 1.0 | 2.2E-03 | 0.9 | 2.7E-05 | 1.4 | 1.2E-09 | 2.9 | 3.1E-07 |
| FmdF | 0.8 | 1.6E-05 | 0.9 | 7.6E-08 | 1.1 | 1.4E-08 | 2.6 | 8.8E-06 |
| FmdG | 1.0 | 3.7E-07 | 0.9 | 1.2E-06 | 1.5 | 3.1E-08 | 2.9 | 1.7E-06 |
| FmdH | 0.7 | 1.5E-06 | 0.4 | 2.1E-06 | 1.2 | 2.0E-05 | 2.4 | 2.3E-06 |
| FdoH | -1.1 | 2.2E-07 | -0.8 | 5.3E-04 | -0.8 | 5.7E-06 | -0.6 | 1.3E-03 |
| FdoG | 0.4 | 3.0E-01 | 0.6 | 8.5E-02 | 1.1 | 2.7E-01 | 1.0 | 1.6E-02 |
| FdoI | NF | NF | NF | NF | NF | NF | NF | NF |
| Gor | 0.2 | 1.8E-06 | 0.2 | 1.4E-04 | 0.2 | 2.2E-05 | 0.2 | 7.4E-06 |
| TtgA | 0.7 | 1.8E-11 | 0.7 | 5.1E-11 | 1.7 | 6.0E-14 | 0.3 | 5.6E-08 |
| TtgB | 0.7 | 6.7E-10 | 0.7 | 7.0E-11 | 1.8 | 1.4E-08 | 0.2 | 1.1E-06 |
| TtgC | 0.6 | 6.2E-09 | 0.7 | 9.2E-12 | 1.7 | 3.7E-15 | 0.2 | 6.4E-08 |
| Ttg2D | -0.2 | 2.6E-02 | 0.1 | 4.1E-02 | 0.1 | 2.0E-02 | 0.0 | 3.5E-01 |
| Ttg2E | -0.1 | 1.3E-02 | 0.0 | 2.6E-03 | -0.3 | 7.5E-06 | -0.2 | 4.6E-06 |
| Ttg2F | 0.1 | 3.9E-02 | 0.0 | 4.7E-02 | -0.2 | 5.7E-01 | -0.1 | 2.5E-01 |

Note: The proteomics data were obtained from four biological replicates. NF: not found.

**Table S4.** Intracellular levels of metabolites (μmol $g_{CDW}^{-1}$) in the central carbon metabolism.

| | SUC | FER | COU | VAN | 4HB |
|---|---|---|---|---|---|
| SUC | -- | 1.94 ± 0.36 | 2.83 ± 0.52 | 1.79 ± 0.21 | 1.50 ± 0.1 |
| Acetyl-CoA | B.D. | 0.11 ± 0.04 | 0.10 ± 0.04 | 0.07 ± 0.03 | 0.07 ± 0.02 |
| Citrate | 0.34 ± 0.09 | 2.88 ± 0.46 | 3.52 ± 0.55 | 1.45 ± 0.20 | 1.02 ± 0.06 |
| α-Ketoglutarate | 0.22 ± 0.08 | 0.98 ± 0.14 | 1.48 ± 0.27 | 1.46 ± 0.19 | 0.75 ± 0.04 |
| Fumarate | 0.63 ± 0.35 | B.D. | B.D. | B.D. | B.D. |
| Malate | 0.53 ± 0.18 | 0.45 ± 0.01 | 0.28 ± 0.09 | 0.15 ± 0.02 | 0.19 ± 0.02 |
| Aspartate | 0.12 ± 0.01 | 3.47 ± 0.30 | 5.13 ± 0.47 | 2.54 ± 0.38 | 4.01 ± 0.40 |
| Pyruvate | 0.65 ± 0.20 | 0.46 ± 0.09 | 0.81 ± 0.07 | 0.65 ± 0.04 | 0.71 ± 0.07 |
| 3PG | 0.20 ± 0.03 | 12.17 ± 0.69 | 8.14 ± 0.92 | 1.50 ± 0.28 | 3.53 ± 0.34 |
| PEP | 0.47 ± 0.15 | 1.93 ± 0.25 | 2.11 ± 0.24 | 1.18 ± 0.23 | 1.89 ± 0.24 |
| G6P | 0.16 ± 0.03 | 0.04 ± 0.02 | 0.10 ± 0.05 | 0.05 ± 0.02 | 0.11 ± 0.01 |
| FBP | 0.24 ± 0.02 | 0.46 ± 0.12 | 0.16 ± 0.06 | 0.17 ± 0.03 | 0.19 ± 0.01 |

Note: Shown values are mean ± standard deviation from four biological replicates. B.D.: Below detection limit. The intracellular levels of metabolites from SUC-grown culture were adapted from Wilkes et al.[1]

**Table S5.** Sum-of-squared residuals (SSR) from models with the activation or deactivation of pyruvate decarboxylation.

| | Activate | Deactivate | Expected SSR range for acceptable fit |
|---|---|---|---|
| FER | 192.7 ± 2.5 | 225.4 ± 2.3 | 139.4-212.4 |
| COU | 157.8 ± 5.7 | 136641.8 ± 3.0 | 142.1-215.7 |

Note: Shown are mean ± standard deviation of SSR values from three independent fits.

**Table S6.** Strains utilized in this study and corresponding construction details.

| Strain | Genotype | Construction details | Reference |
|---|---|---|---|
| *P. putida* | Wild-type *Pseudomonas putida* KT2440 | n/a | ATCC® 47054 |
| AG5577 | *P. putida* KT2440 ΔPP_4740::BxB1_RV_phi370_attB cassette ΔPP_4217/4218 intergenic::TG1_BL3_A118_attB cassette ΔPP_2876::R4_phiBT1_MR11_attB cassette | Contains a total of nine attB attachment sites (3 in each of 3 loci) for use with serine integrase helper plasmids. See reference for more details. | Schmidt et al., 2023 [4] |
| RW123 | *P. putida* KT2440 ΔPP_4740::BxB1attL:**$P_{tac}$:*pcaHG***:attR ΔPP_4740::RV_phi370_attB cassette ΔPP_4217/4218 intergenic::TG1_BL3_A118_attB cassette ΔPP_2876::R4_phiBT1_MR11_attB cassette | pRW44 was transformed into AG5577 using SAGE integration with helper plasmid pGW31. Integration of **$P_{tac}$:*pcaHG*** was confirmed by colony PCR with oQP1357 and oQP1358 (Tm=67°C) followed by Oxford Nanopore sequencing at Plasmidsaurus. | This study |
| RW124 | *P. putida* KT2440 ΔPP_4740::BxB1_RV_phi370_attB cassette ΔPP_4217/4218 intergenic::TG1-attL:**$P_{tac}$:*vanAB***:attR ΔPP_4217/4218 intergenic::BL3_A118_attB cassette ΔPP_2876::R4_phiBT1_MR11_attB cassette | pRW46 was transformed into AG5577 using SAGE integration with helper plasmid pGW38. Integration of **$P_{tac}$:*vanAB*** was confirmed by colony PCR with oJE529 and oJE1628 (Tm=67°C) followed by Oxford Nanopore sequencing at Plasmidsaurus. | This study |
| RW125 | *P. putida* KT2440 ΔPP_4740::BxB1attL:**$P_{tac}$:*pcaHG***:attR ΔPP_4740::RV_phi370_attB cassette ΔPP_4217/4218 intergenic::TG1-attL:**$P_{tac}$:*vanAB***:attR ΔPP_4217/4218 intergenic::BL3_A118_attB cassette ΔPP_2876::R4_phiBT1_MR11_attB cassette | pRW46 and pRW44 were transformed into AG5577 using SAGE integration with helper plasmids pGW31 and pGW38. Integration of **$P_{tac}$:*pcaHG*** was confirmed by colony PCR with oQP1357 and oQP1358 (Tm=67°C, 7 kB) and integration of **$P_{tac}$:*vanAB*** was confirmed by colony PCR with oJE529 and oJE1628 (Tm=67°C). Both PCR products were sequence confirmed using Oxford Nanopore sequencing at Plasmidsaurus | This study |
| RW126 | *P. putida* KT2440 ΔPP_4740::BxB1attL:**$P_{tac}$:*pcaHG***:attR ΔPP_4740::RV_phi370_attB cassette ΔPP_4217/4218 intergenic::TG1-attL:**$P_{tac}$:*vdh*:*vanAB***:attR ΔPP_4217/4218 intergenic::BL3_A118_attB cassette ΔPP_2876::R4_phiBT1_MR11_attB cassette | pRW48 and pRW44 were transformed into AG5577 using SAGE integration with helper plasmids pGW31 and pGW38. Integration of **$P_{tac}$:*pcaHG*** was confirmed by colony PCR with oQP1357 and oQP1358 (Tm=67°C, 7 kB) and integration of **$P_{tac}$:*vdh*:*vanAB*** was confirmed by colony PCR with oJE529 and oJE1628 (Tm=67°C). Both PCR products were sequence confirmed using Oxford Nanopore sequencing at Plasmidsaurus | This study |

| | | | |
|---|---|---|---|
| RW127 | *P. putida* KT2440 ΔPP_4740::BxB1_RV_phi370_attB cassette ΔPP_4217/4218 intergenic::TG1-attL:**$P_{tac}$:*vdh*:**attR ΔPP_4217/4218 intergenic::BL3_A118_attB cassette ΔPP_2876::R4_phiBT1_MR11_attB cassette | pRW45 was transformed into AG5577 using SAGE integration with helper plasmid pGW38. Integration of **$P_{tac}$:*vdh*** was confirmed by colony PCR with oJE529 and oJE1628 (Tm=67°C) followed by Oxford Nanopore sequencing at Plasmidsaurus. | This study |
| RW128 | *P. putida* KT2440 ΔPP_4740::BxB1_RV_phi370_attB cassette ΔPP_4217/4218 intergenic::TG1-attL:**$P_{tac}$:*pobA*:**attR ΔPP_4217/4218 intergenic::BL3_A118_attB cassette ΔPP_2876::R4_phiBT1_MR11_attB cassette | pRW47 was transformed into AG5577 using SAGE integration with helper plasmid pGW38. Integration of **$P_{tac}$:*pobA*** was confirmed by colony PCR with oJE529 and oJE1628 (Tm=67°C) followed by Oxford Nanopore sequencing at Plasmidsaurus. | This study |
| RW129 | *P. putida* KT2440 ΔPP_4740::BxB1attL:**$P_{tac}$:*pcaHG***:attR ΔPP_4740::RV_phi370_attB cassette ΔPP_4217/4218 intergenic::TG1-attL:**$P_{tac}$:*pobA*:**attR ΔPP_4217/4218 intergenic::BL3_A118_attB cassette ΔPP_2876::R4_phiBT1_MR11_attB cassette | pRW47 and pRW44 were transformed into AG5577 using SAGE integration with helper plasmids pGW31 and pGW38. Integration of **$P_{tac}$:*pcaHG*** was confirmed by colony PCR with oQP1357 and oQP1358 (Tm=67°C, 7 kB) and integration of **$P_{tac}$:*pobA*** was confirmed by colony PCR with oJE529 and oJE1628 (Tm=67°C). Both PCR products were sequence confirmed using Oxford Nanopore sequencing at Plasmidsaurus. | This study |

**Table S7.** Oligonucleotides utilized in this study.

| Name | Sequence (5'->3') | Purpose |
|---|---|---|
| oQP1357 | GTCTTTATAGCTTCGACGTCATAGTAGG | For verification of integration of BxB1 site. |
| oQP1358 | GAACAGCAGGTTGGTCTTGATGC | |
| oJE529 | CGGCACTTCGCCCAATA | For verification of integration at TG1 site. |
| oJE1628 | ACGCCTGCTTCATTGAACTT | |

**Table S8.** Plasmids utilized in this study.

| Name | Description | Construction details | Reference |
|---|---|---|---|
| pRW44 | Integrates $P_{tac}$:*pcaHG* at the BxB1 site. | Synthesized and cloned into pJH419 vector by Twist Biosciences. Sequence of insert includes $P_{tac}$ promoter, ribosome binding site, and native *pcaHG* from KT2440. | This study |
| pRW45 | Integrates $P_{tac}$:*vdh* at the TG1 site. | Synthesized and cloned into pJH210 vector by Twist Biosciences. Sequence of insert includes $P_{tac}$ promoter, ribosome binding site, and native *vdh* from KT2440. | This study |
| pRW46 | Integrates $P_{tac}$:*vanAB* at the TG1 site. | Synthesized and cloned into pJH210 vector by Twist Biosciences. Sequence of insert includes $P_{tac}$ promoter, ribosome binding site, and native *vanAB* from KT2440. | This study |
| pRW47 | Integrates $P_{tac}$:*pobA* at the TG1 site. | Synthesized and cloned into pJH210 vector by Twist Biosciences. Sequence of insert includes $P_{tac}$ promoter, ribosome binding site, and native *pobA* from KT2440. | This study |
| pRW48 | Integrates $P_{tac}$:*vdh:vanAB* at the TG1 site. | Synthesized and cloned into pJH210 vector by Twist Biosciences. Sequence of insert includes $P_{tac}$ promoter, ribosome binding site, and native *vdh:vanAB* from KT2440. | This study |
| pJH419 | SAGE cargo target/attP plasmid. Contains attP sites for the BxB1 serine integrase. ColE1 origin, spectinomycin/streptomycin and ampicillin resistance markers. | Donated from the laboratory of Adam M. Guss | Elmore et al., 2023[6] |
| pJH210 | SAGE cargo target/attP plasmid. Contains attP sites for the TG1 serine integrase. ColE1 origin, kanamycin resistance marker. | Donated from the laboratory of Adam M. Guss | Elmore et al., 2023[6] |
| pGW31 | Helper plasmid ColE1 vector with apramycin marker expressing BxB1 integrase under the control of $P_{tac}$ | Donated from the laboratory of Adam M. Guss | Elmore et al., 2023[6] |
| pGW38 | Helper plasmid ColE1 vector with apramycin marker expressing TG1 integrase under the control of $P_{tac}$ | Donated from the laboratory of Adam M. Guss | Elmore et al., 2023[6] |
